# Development of a soil inoculation method coupled with blocker-mediated 16S rRNA gene amplicon sequencing reveals the effect of antibacterial T6SS on agrobacteria tumorigenesis and gallobiome composition

**DOI:** 10.1101/2022.10.28.514271

**Authors:** Si-Chong Wang, Ai-Ping Chen, Shu-Jen Chou, Chih-Horng Kuo, Erh-Min Lai

## Abstract

The type VI secretion system (T6SS) is deployed by many proteobacteria to secrete effector proteins into bacterial competitors for competition or eukaryotic cells for pathogenesis. Agrobacteria, a group of soil-borne phytopathogens causing crown gall disease on various plant species, deploys the T6SS to attack closely- and distantly-related bacterial species *in vitro* and *in planta*. Current evidence suggests that the T6SS is not essential for pathogenesis under direct inoculation but it remains unknown whether the T6SS influences natural disease incidence or the microbial community within crown galls (i.e., gallobiome). To address these two key questions, we established a soil inoculation method on wounded tomato seedlings that mimics natural infections and developed a bacterial 16S rRNA gene amplicon enrichment sequencing platform. By comparing the *Agrobacterium* wild-type strain C58 with two T6SS mutants, we demonstrate that the T6SS influences both disease occurrence and gallobiome composition. Based on multiple inoculation trials across seasons, all three strains could induce tumors but the mutants had significantly lower disease incidences. The season of inoculation played a more important role than the T6SS in shaping the gallobiome. The influence of T6SS was evident in summer, in which two *Sphingomonas* species and the family Burkhoderiaceae were enriched in the gallobiome induced by the mutants. Further *in vitro* competition and colonization assay demonstrated the T6SS-mediated antagonism to a *Sphingomonas* sp. R1 strain isolated from tomato rhizosphere in this study. In conclusion, this work demonstrates that the *Agrobacterium* T6SS promotes tumorigenesis in infection process and provides competitive advantages in gall-associated microbiota.

**IMPORTANCE:** The T6SS is widespread among Proteobacteria and used for interbacterial competition by agrobacteria, which are soil inhabitants and opportunistic bacterial pathogens causing crown gall disease in a wide range of plants. Current evidence indicates that the T6SS is not required for gall formation when agrobacteria are inoculated directly on plant wounding sites. However, in natural settings, agrobacteria may need to compete with other bacteria in bulk soil to gain access to plant wounds and influence microbial community inside crown galls. The role of the T6SS in these critical aspects of disease ecology have remained largely unknown. In this study, we successfully developed a <u>S</u>oil <u>I</u>noculation method coupled with <u>B</u>locker-mediated enrichment of <u>Bac</u>terial 16S rRNA gene Amplicon <u>Seq</u>uencing, named as SI-BBacSeq, to address these two important questions. We provided evidence that the T6SS promotes disease occurrence and influences crown gall microbiota composition by interbacterial competition.

## INTRODUCTION

Many proteobacteria, including pathogens and commensals, deploy the type VI secretion system (T6SS) for antagonism or pathogenesis (1, 2). The T6SS is a protein translocation apparatus used to inject effectors into target cells, mainly in a contact-dependent manner. Based on the destination and biological function of known effectors (3), the T6SS mainly functions as an antibacterial weapon used by bacteria to inhibit or kill the competing bacterial species, thus providing a competitive advantage and shaping microbiota in their ecological niche (4).

Previous studies on the microbiome associated with animal guts or plants indicated that the T6SS genes are enriched in these communities, suggesting T6SS may be important for niche competition (5–8). Metagenomic analysis of human gut microbiota revealed a role of the T6SS for the domination of the gut symbiont *Bacteroides fragilis* by targeting other members of microbiome *in vitro* and *in vivo* (6, 7). A recent study further showed that a murine pathogen *Citrobacter rodentium* and resident commensal Enterobacteriaceae share the same strategy by using the T6SS for niche competition in the murine gastrointestinal tract (9). The T6SS is also deployed by plant pathogens to gain competitive growth advantage *in planta* as well as for beneficial bacteria to prevent or reduce disease symptoms caused by competing pathogens (10–12). Comparative metagenomic analysis of microbiota between T6SS^+^ and T6SS^-^ bacterial strains were also carried out. The gut microbiota of pests infected by *Pseudomonas protegens*, a plant-beneficial bacterium capable of invading insect pests, showed that the T6SS has no significant impact on microbiota diversity at phylum/class level but affects the abundance of Enterobacteriaceae (13). These studies collectively suggest that the T6SS is a potent antibacterial weapon used by invading pathogens or resident bacteria to gain competitive advantage in their ecological niches. However, the knowledge regarding the degree of the T6SS in shaping microbiota and the molecular mechanisms of interbacterial competition in complex microbial community are limited.

Agrobacteria are a diverse group of bacteria that include members from several genera (14). These plant pathogens are capable of inducing crown gall or hairy root disease on plants by transferring a piece of DNA named T-DNA from bacteria into plants via the type IV secretion system (T4SS) (15). The T6SS is highly conserved in several *Agrobacterium* species and plays a role in interbacterial competition (16–18). Among these *Agrobacterium* species, the T6SS is encoded by a gene cluster consisting of an *imp* operon encoding the main T6SS components and an *hcp* operon encoding the puncturing device and effectors (10, 18, 19). Using a key *Agrobacterium* reference strain C58, which is commonly known as a member of *A. tumefaciens* but recently reclassified as *A. fabrum* (20), we previously discovered that it uses a T6SS DNase effector to gain competitive growth advantage *in vitro* and *in planta* (10). Interestingly, agrobacteria with incompatible effector-immunity (EI) pairs exhibit strong antagonism between species level, while only weak or non-detectable effect within species level (16). Moreover, higher T6SS-mediated killing outcome was observed when nutrient is scarce, as opposed to nutrient rich condition (21). Thus, genetic and environmental factors beyond EI pairs also contributes to interbacterial competition.

To date, agrobacterial T6SS has been only demonstrated as an antibacterial weapon (16–18). No evidence for its role in promoting virulence when tumor assays were conducted in sterile condition or directly inoculated on the stems of various plant species including tomato plants (19). Considering that agrobacteria may need to compete with other bacteria in bulk soil or rhizosphere to gain access to plant wounds for inciting crown galls, we reasoned that agrobacterial T6SS may influence microbial community and pathogenesis under a more natural setting. To address these questions, we developed a soil inoculation protocol to mimic natural infection on wounded tomato seedlings across seasons for evaluating the impact of agrobacterial T6SS on disease occurrence and crown gall microbiota (termed gallobiome). Moreover, to overcome the challenge of interference of host organelles in the study of plant-associated microbiota, we optimized a blocker-based method for enriching true bacterial reads in 16S rRNA gene amplicon sequencing. Based on results from our infection assays, gallobiome composition analysis, as well as *in vitro* competition and colonization assays, this work indicates that agrobacterial T6SS may provide competitive advantages on plant surface for effective infection, leading to a higher disease incidence.

## RESULTS

### Environmental factors and T6SS affect crown gall disease incidence

The wild type (WT) strain C58 and two C58-derived T6SS mutants with deletion of essential T6SS genes, Δ*tssL* and Δ*tssB* respectively, were used to study the effects of agrobacterial T6SS in tumorigenesis using a soil inoculation method. Nine batches of inoculation experiments across different seasons were conducted in this study (Table 1). In total, 139 crown galls were collected, including 70 induced by WT, 41 induced by Δ*tssL*, and 28 induced by Δ*tssB*. The results showed that all three strains are capable of inducing tumors, but the disease incidences of Δ*tssL* and Δ*tssB* were significantly lower than WT (Table 1, Fig. 1A). There was no significant weight difference among the crown galls induced by different strains (Fig. 1B). It is notable that there is an inverse correlation between disease incidence and temperature across seasons for all three strains throughout the year (Fig. 1C, 1D). These results indicated that the presence of a functional T6SS and the month of inoculation both affected the disease incidence under soil inoculation condition.

**Figure 1.**
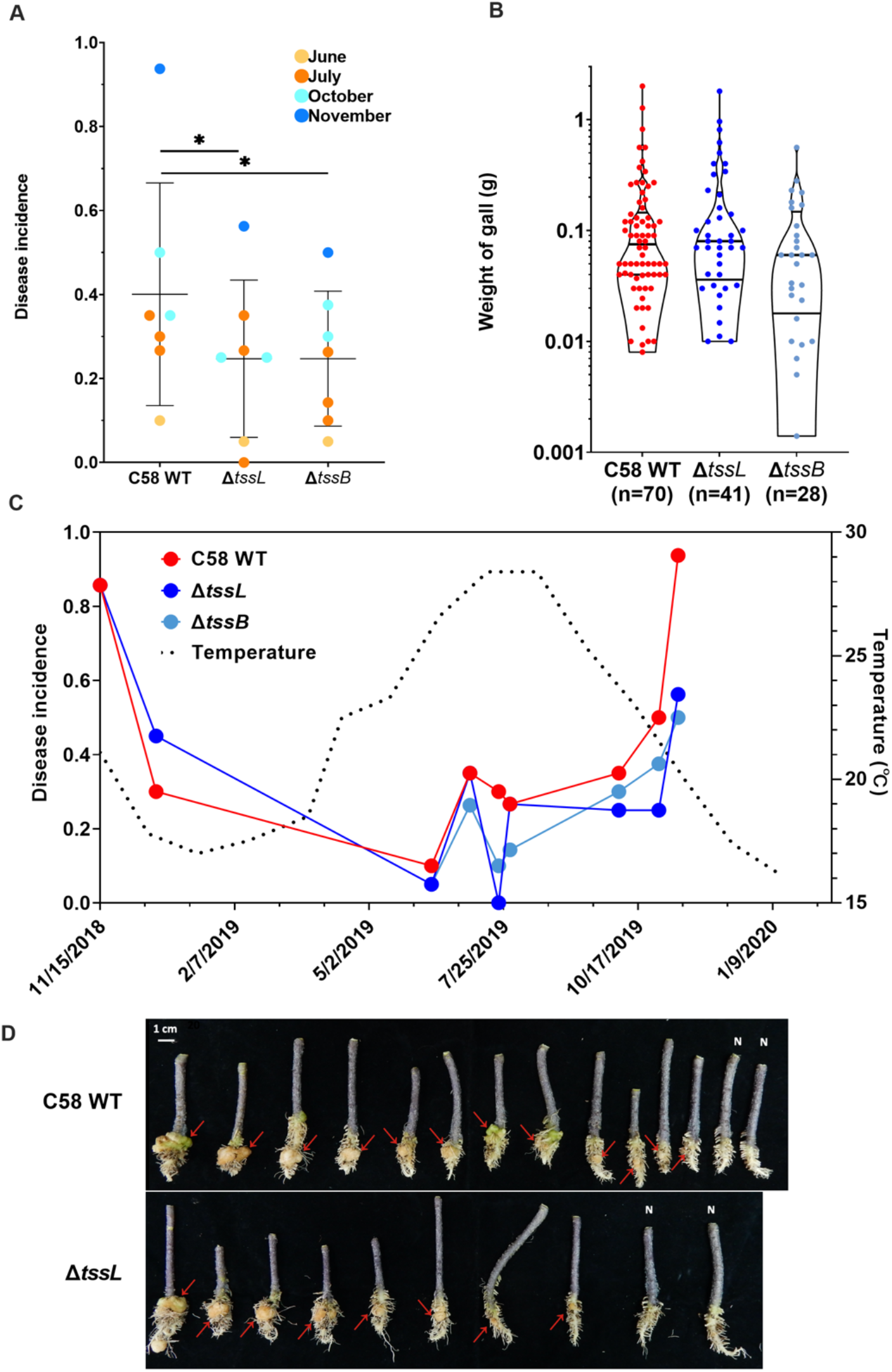
Tumorigenesis assay of agrobacterial strain C58 and its T6SS-deficienct mutants. For each experiment, 14-20 wounded tomato seedlings were grown in soil containing one of the strains tested (i.e., C58 WT, Δ*tssL*, and Δ*tssB*) and collected at 60 days post inoculation (dpi). (A) Disease incidence plotted by strain and color-coded according to the month of inoculation (light yellow, June; orange, July; light blue, October; dark blue, November). Lines and error bars indicate mean ± SD. Statistical significance was tested using two-way ANOVA followed by Tukey’s multiple comparisons; *p*-value = 0.025 and 0.027 when comparing the wild type to Δ*tssL* and Δ*tssB*,respectively. (B) Weight distribution of the galls collected (See details in Table S7). No significant difference among the three strains (*p*-value= 0.15, Kruskai-Wallis test). (C) Correlation plot of disease incidence and temperature of daily average. (D) Crown galls generated through soil inoculation, example from inoculation on 11/15/2018. N indicated no gall formation in some of inoculated plants.

**Table 1.** Incidences of crown gall disease on wounded tomato stems by soil inoculation

| Batch | Date of inoculation | Disease incidence (diseased/total inoculated seedlings) |  |  |
| --- | --- | --- | --- | --- |
| | | C58 WT<sup>a</sup> | $\Delta tssL$<sup>b</sup> | $\Delta tssB$<sup>b</sup> |
| 1 | 2018/11/15 | 0.86 (12/14) | 0.86 (12/14) | N.D.* |
| 2 | 2018/12/20 | 0.30 (6/20) | 0.45 (9/20) | N.D. |
| 3 | 2019/06/10 | 0.10 (2/20) | 0.05 (1/20) | 0.05 (1/20) |
| 4 | 2019/07/04 | 0.35 (7/20) | 0.35 (7/20) | 0.26 (5/19) |
| 5 | 2019/07/22 | 0.30 (6/20) | 0.00 (0/20) | 0.10 (2/20) |
| 6 | 2019/07/29 | 0.27 (4/15) | 0.27 (4/15) | 0.14 (2/14) |
| 7 | 2019/10/05 | 0.35 (7/20) | 0.25 (5/20) | 0.30 (6/20) |
| 8 | 2019/10/30 | 0.50 (8/16) | 0.25 (4/16) | 0.38 (6/16) |
| 9 | 2020/11/11 | 0.94 (15/16) | 0.56 (9/16) | 0.50 (8/16) |

<sup>a, b</sup>: Two-way ANOVA followed by Tukey's multiple comparisons test on disease incidences of different strains indicated the disease incidence of $\Delta tssL$ and $\Delta tssB$ was significantly lower than *Agrobacterium* C58 WT (*p*-value are 0.0225 and 0.0227 respectively). \*: Not Determined.

### Round I of 16S rRNA gene amplicon sequencing: Initial trial

To determine the gallobiome composition, crown galls with similar weights across seasons were selected for 16S rRNA gene amplicon sequencing. For Round I, six crown gall DNA samples, three induced by WT and three by Δ*tssL* were amplified with two commonly used 16S rRNA gene primer sets V3-4 and V5-7 (Table S1). The Illumina sequencing produced 140,830 and 601,299 reads from V3-4 and V5-7 sets, respectively. However, the majority (99.1% in V3-4 and 93.6% for V5-7) of these reads were derived from plant chloroplast and mitochondria. After removing these host contaminations, only 1,230 and 38,320 bacterial reads remained (Table 2).

**Table 2.** Sum of read counts in six tumor samples before and after filtering non-bacterial reads from amplicon sequencing Run I for protocol test

| 16S rRNA regions | Before filtering | Bacterial reads | Percentage of non-bacterial DNA |
| --- | --- | --- | --- |
| V3-4 | 140,830 | 1,230 | 99.1% |
| V5-7 | 601,299 | 38,320 | 93.6% |

### Round II of 16S rRNA gene amplicon sequencing: Optimization

Due to the severe host contamination observed in Round I that used the universal 16S rRNA gene primers, we evaluated the performance of different 16S rRNA gene primers with blockers (3’ modified oligonucleotides with C3 spacer) that could prevent the amplification of tomato chloroplast and mitochondrial rRNA genes (22–24).

Based on the alignment of bacterial, tomato chloroplast and mitochondrial 16S rRNA gene sequences (Fig. S1), we designed the primer sets and cognate blockers against different variable region in 16S rRNA genes (Table S1 and S2). The new primer sets and blockers were tested using the three WT-induced crown gall DNA samples from Round I. After sequencing and data processing, the result indicated that the blockers have variable effectiveness in reducing host contaminations (Table 3 and Fig. 2A). For V3-4 and V5-7 primer sets, adding blockers increased the bacterial reads to 5.3-41.5% of total reads, compared to only 0.2-1.9% of bacterial reads per sample without blockers. For V1-3 and V6-8 primer sets, adding blockers only increased the bacterial reads to 1.6-5.6% of total reads per sample. The reads were grouped into species-level operational taxonomic units (OTUs) with 99% sequence identity. The alpha rarefaction curves based on the OTU counts indicated that, for those crown gall samples, the curves would approach saturation when the sequencing depth was over about 10,000 reads per sample (Fig. 2B). According to the bar plot of bacterial composition at family levels, we could observe higher number of families identified from the same crown galls amplified by adding blockers (Fig. 3A). The result of Principal Coordinate Analysis (PCoA) indicated that adding blockers did not cause significant biases in the inferred microbiota composition (Fig. 3B). However, the use of different primer sets resulted in significant difference in the inferred microbiota composition (Fig. 3C), which has been reported previously (25). There is also significant difference among different crown galls (Fig. 3D).

**Figure 2.**
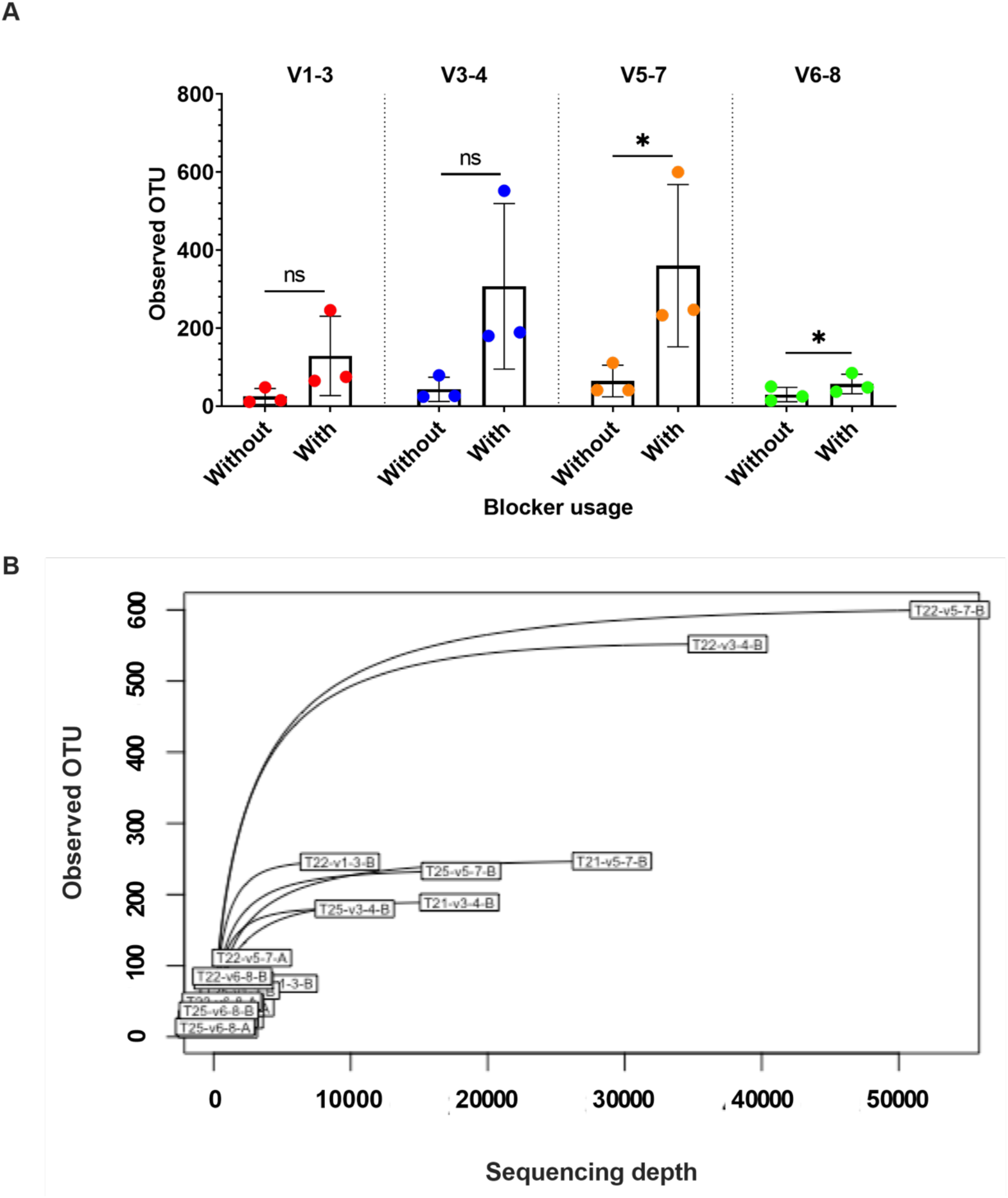
Optimizing 16S rRNA gene amplification by using PCR blockers. Three crown gall samples (named as 21W, 22W and 25W) induced by C58 wild type were used for analysis. V1-3, V3-4, V5-7 and V6-8 represent different primer pairs targeting different variable regions on 16S rRNA gene. (A) The OTU counts of 16 rRNA gene amplicons after filtering non-bacterial OTUs. (ns, not statistically significant; *, *p-value* < 0.05, student’s t-test). The bar indicated mean of OTUs ± SD. (B) Alpha rarefaction curves of the observed bacterial OTUs based on datasets from amplicon sequencing run II. Sample size in X-axis indicates different sub-sampling depth of each dataset.

**Figure 3.**
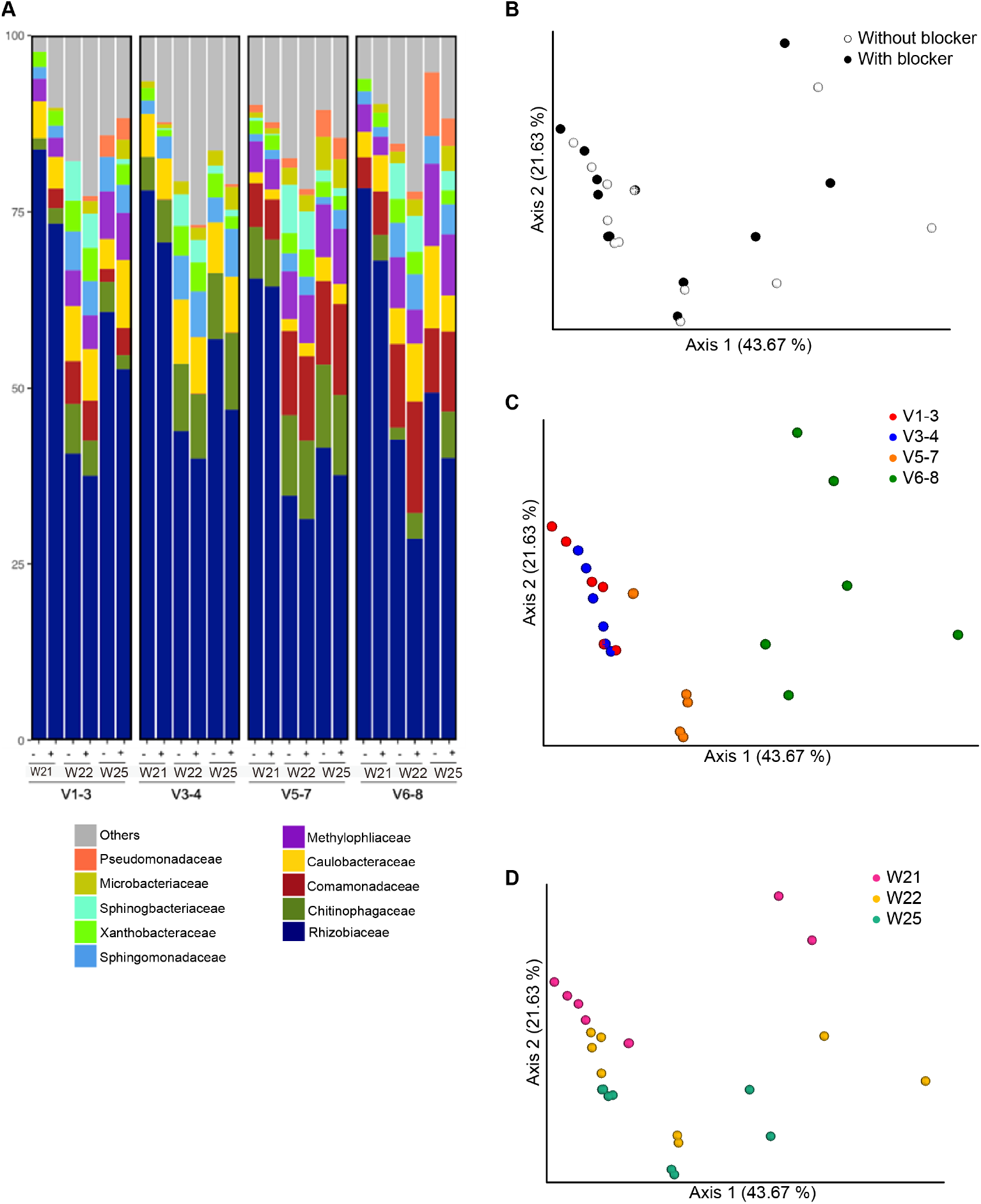
Gallobiome composition when using different primers amplified with or without blockers. (A) Different primer sets (V1-3, V3-4, V5-7 and V6-8)for analysis of three crown gall samples (21W, 22W and 25W, see details in Table S8) induced by C58 wild type were indicated. Top 10 observed families were listed. “-”: amplification without blockers; “+”: amplification with blockers. Principle coordinate analysis (PCoA) plots of the bacterial composition of the datasets were generated based on the weighted Unifrac matrix, and color coded by (B) blocker usage (*p-*value = 0.068, R^2^ = 0.026, ADONIS), (C) different primer sets (*p*-value = 0.001, R^2^ = 0.524, ADONIS) and (D) different crown gall samples (*p*-value = 0.001, R^2^ = 0.238, ADONIS).

**Table 3.** The read count before and after filtering non-bacterial ASVs from the three crown gall DNA samples (named as W21, W22, and W25) induced by *Agrobacterium* C58 wild type

| Sample-id | Dataset-id | Read before filtering | Reads after filtering chloroplast and mitochondrial DNA | Percentage of non-host reads |
| --- | --- | --- | --- | --- |
| 1115-21W | T21-v1-3-B | 176,544 | 9,914 | 5.6% |
| 1115-22W | T22-v1-3-B | 183,475 | 5,608 | 3.1% |
| 1115-25W | T25-v1-3-B | 152,387 | 2,404 | 1.6% |
| 1115-21W | T21-v1-3-A | 185,107 | 853 | 0.5% |
| 1115-22W | T22-v1-3-A | 247,240 | 379 | 0.2% |
| 1115-25W | T25-v1-3-A | 178,980 | 220 | 0.1% |
| 1115-21W | T21-v3-4-B | 239,243 | 38,462 | 16.1% |
| 1115-22W | T22-v3-4-B | 246,063 | 19,218 | 7.8% |
| 1115-25W | T25-v3-4-B | 209,991 | 11,185 | 5.3% |
| 1115-21W | T21-v3-4-A | 200,185 | 979 | 0.5% |
| 1115-22W | T22-v3-4-A | 205,966 | 796 | 0.4% |
| 1115-25W | T25-v3-4-A | 193,350 | 426 | 0.2% |
| 1115-21W | T21-v5-7-B | 129,422 | 53,715 | 41.5% |
| 1115-22W | T22-v5-7-B | 140,012 | 29,078 | 20.8% |
| 1115-25W | T25-v5-7-B | 135,692 | 18,052 | 13.3% |
| 1115-21W | T21-v5-7-A | 150,015 | 2,792 | 1.9% |
| 1115-22W | T22-v5-7-A | 139,974 | 1,507 | 1.1% |
| 1115-25W | T25-v5-7-A | 133,035 | 786 | 0.6% |
| 1115-21W | T21-v6-8-B | 92,167 | 3,691 | 4.0% |
| 1115-22W | T22-v6-8-B | 92,053 | 3,197 | 3.5% |
| 1115-25W | T25-v6-8-B | 82,361 | 2,077 | 2.5% |
| 1115-21W | T21-v6-8-A | 121,145 | 1,604 | 1.3% |
| 1115-22W | T22-v6-8-A | 138,186 | 1,273 | 0.9% |
| 1115-25W | T25-v6-8-A | 121,820 | 708 | 0.6% |
|  | NC-v3-4-B | 1,012 | 979 | 96.7% |

Overall, adding blockers at the PCR step increased bacterial read counts and the resolution of gallobiome. Because the use of V5-7 primers and cognate blockers obtained the highest number and percentage of bacterial reads, V5-7 set-up with blockers were selected for 16 rRNA gene amplicon sequencing of crown galls induced by WT and T6SS mutants.

### Round III of 16S rRNA gene amplicon sequencing: Impact of the T6SS on gallobiome

Among all of 139 crown galls collected, 53 tumors in the range of 0.06 – 0.56 g (24 induced by WT, 16 by Δ*tssL*, and 13 by Δ*tssB*) were used for the analysis. We obtained an average of 6,9980 ± 26,452 (mean ± SD) reads per sample, ranged from 9,814 to 116,072 reads per sample. Before the analysis, the amplicon sequence variants (ASVs) were clustered into species-level OTUs and the singletons were removed. Based on the alpha rarefaction curves and the minimal read counts of the samples (Fig. S2), diversity analysis was conducted with datasets subsampling at 9800 reads per sample. The result of PCoA showed that the gallobiome induced by WT and two T6SS mutant strains were not significantly different (Fig. 4A). Most of the variation between samples were contributed by different seasons (July vs. October/November) (Fig. 4B). After splitting and re-analyzing the dataset based on the month of inoculation, we observed the difference between the gallobiomes associated with WT and two T6SS mutants in July (Fig. 4C). In contrast, no difference was detected in those galls induced in October or November (Fig. 4D and 4E).

**Figure 4.**
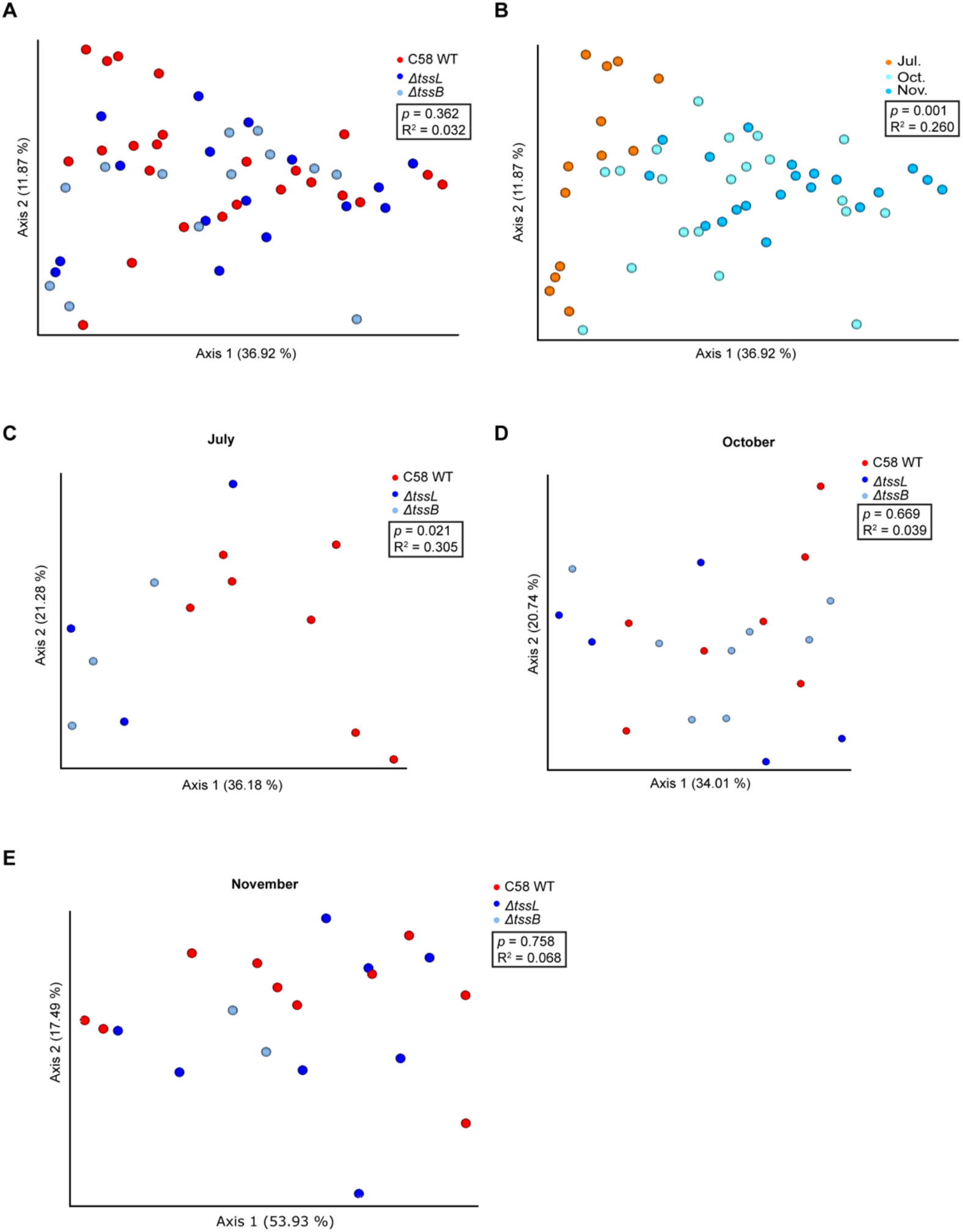
Principle coordinate analysis (PCoA) of bacterial composition in 53 crown galls induced by C58 WT and T6SS mutants. The plots were drawn based on weighted Unifrac matrix of the bacterial communities. Crown galls induced by different strains (C58 WT, Δ*tssB*, Δ*tssL*) (A) or inoculated in different months (B) were labeled with different colors in each panel. The datasets were further split based on the month of inoculation, and the PCoA plots of gallobiomes from (C) July, (D) October and (E) November were indicated above. Statistical differences in clustering were evaluated via ADONIS permutation test, and the corresponding *p*-values and R^2^ are indicated.

The alpha diversity indices including observed OTUs, Shannon index, and Pielou’s evenness index had no significant difference between the gallobiomes associated with WT and T6SS mutants (Fig. 5A, 5B and 5C). Interestingly, we found that, in galls induced in July, the WT gallobiome showed significantly higher Faith’s phylogenic diversity than the gallobiome associated with Δ*tssL*; and in galls induced in November, the Δ*tssL* gallobiome showed higher diversity than the WT gallobiome (Fig. 5D). The gallobiome associated with Δ*tssB* also exhibited higher diversity than the WT gallobiome but the difference was not statistically significant.

**Figure 5.**
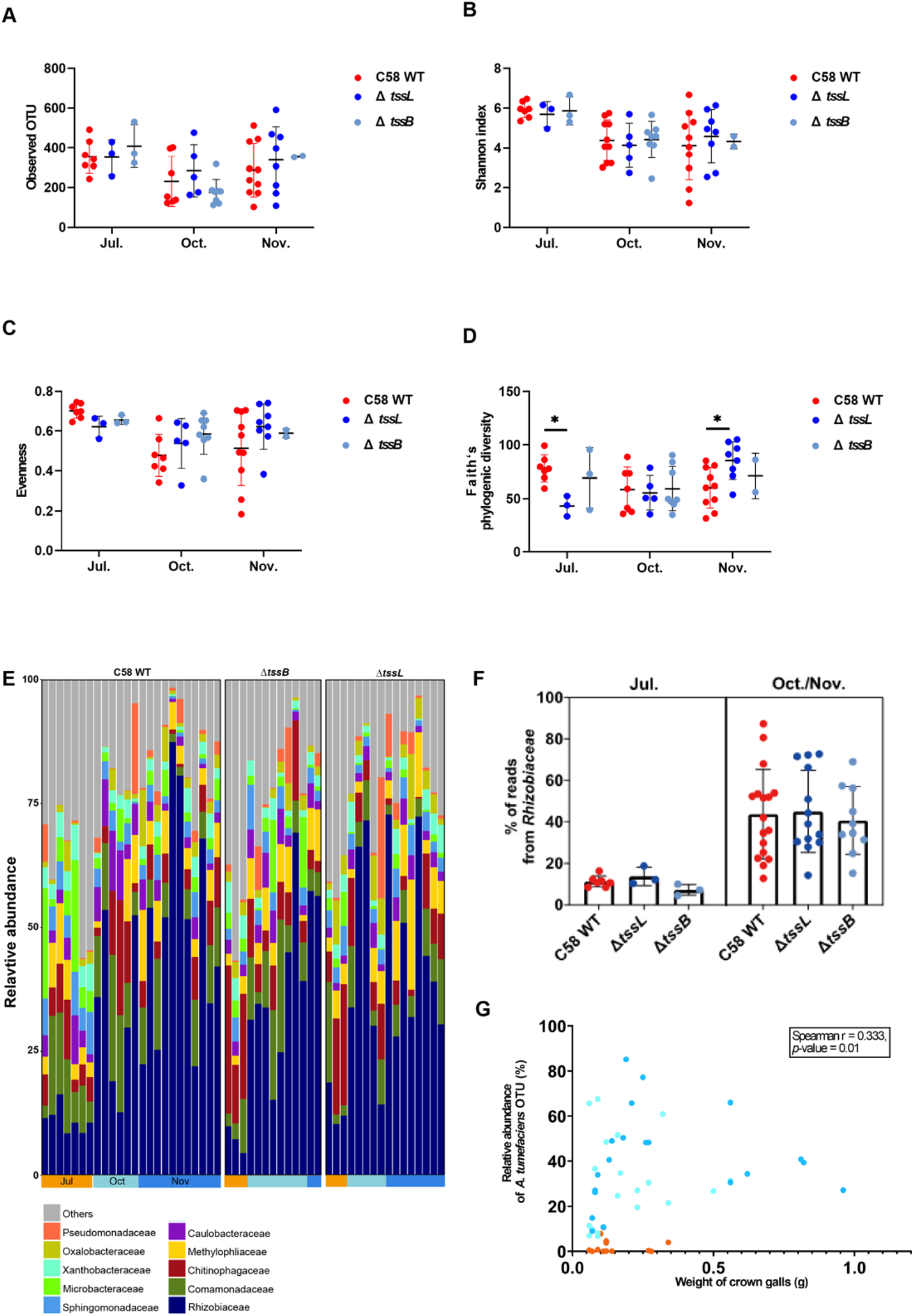
Alpha diversity and composition of crown galls generated in different months by C58 WT and T6SS mutants. (A) Observed OTU, (B) Shannon index, (C) evenness, and (D) Faith’s phylogenic diversity of each sample generated in different months was indicated. Lines and error bars indicate mean ± SD. Two-way ANOVA followed by Tukey’s multiple comparisons test were performed, and the asterisks indicated the statistically significant difference between two strains (*p*-value < 0.05). (E) Bacterial composition of 53 crown galls. For each strain, the samples are grouped by the month of inoculation. Top-ten most abundant families were listed and colored differently, and the remaining ones are combined into “Others”. (F) Relative abundance of Rhizobiaceae in galls inoculated in different months and strains. (G) Scatter plot of relative abundance of agrobacteria OTU in gallobiome - weight of the crown galls. A positive correlation between weight of crown gall and relative abundance of *Agrobacterium* C58 was observed (Spearman r = 0.333, *p*-value = 0.01).

While there were variations in bacterial compositions among crown galls, the top 10 bacterial families in gallobiomes were quite consistent, including: Rhizobiaceae, Comamonadaceae, Chitinophagaceae, Methyphliaceae, Caulobacteraceae, Sphingomonadaceae, Microbacreriaceae, Xanthobacteraceae, Oxalobacteraceae, and Pseudomonaceae (Fig. 5E). The bacterial family Rhizobiaceae, which *Agrobacterium* belongs to, was found to be highly variable and accounted for 5-85% of the entire community. The most abundant OTU in this Rhizobiaceae dataset was 100% matched to 16S rRNA genes of C58, suggesting that the abundance of this OTU could be referred to the relative abundance of WT or mutant inoculum in crown galls. When the weight of crown galls was plotted against the relative abundance of this OTU in gallobiomes, we found that the abundance of Rhizobiaceae in July was dramatically lower than those in October and November (Fig. 5F), but there was no significant difference observed among different strains. Furthermore, a positive correlation between weight of crown galls and relative abundance of this agrobacterial OTU was observed (Fig. 5G).

### Sphingomonadaceae and Burkholderiacea were more abundant in the gallobiomes induced by the T6SS mutants in July

Next, we analyzed the species-level OTU and family with differential abundance (DA) between gallobiome associated with WT and T6SS mutants in July. Two OTUs belonged to Sphingomonadaceae, named as SphinOTU1 and SphinOTU2 here, and Burkhoderiaceae family were significantly enriched in the gallobiomes induced by T6SS mutants (Fig. 6A). SphinOTU1 was only present in gallobiomes in July but not in October/November, and SpinOTU2 was present in both July and October but not in November (Fig 6B). Neither SpinOTU1 or SpinOTU2 was identified in gallobiomes induced by the WT in July. Burkhoderiaceae was enriched in gallobiomes induced by the T6SS mutants in July, but no consistent enrichment in October and November (Fig 6C). Although SphinOTU1 and SphinOTU2 were only present in gallobiomes induced by T6SS mutants in July; at family level, Sphingomonadaceae was not enriched in T6SS-dependent manner in any months (Fig. 6C).

**Figure 6.**
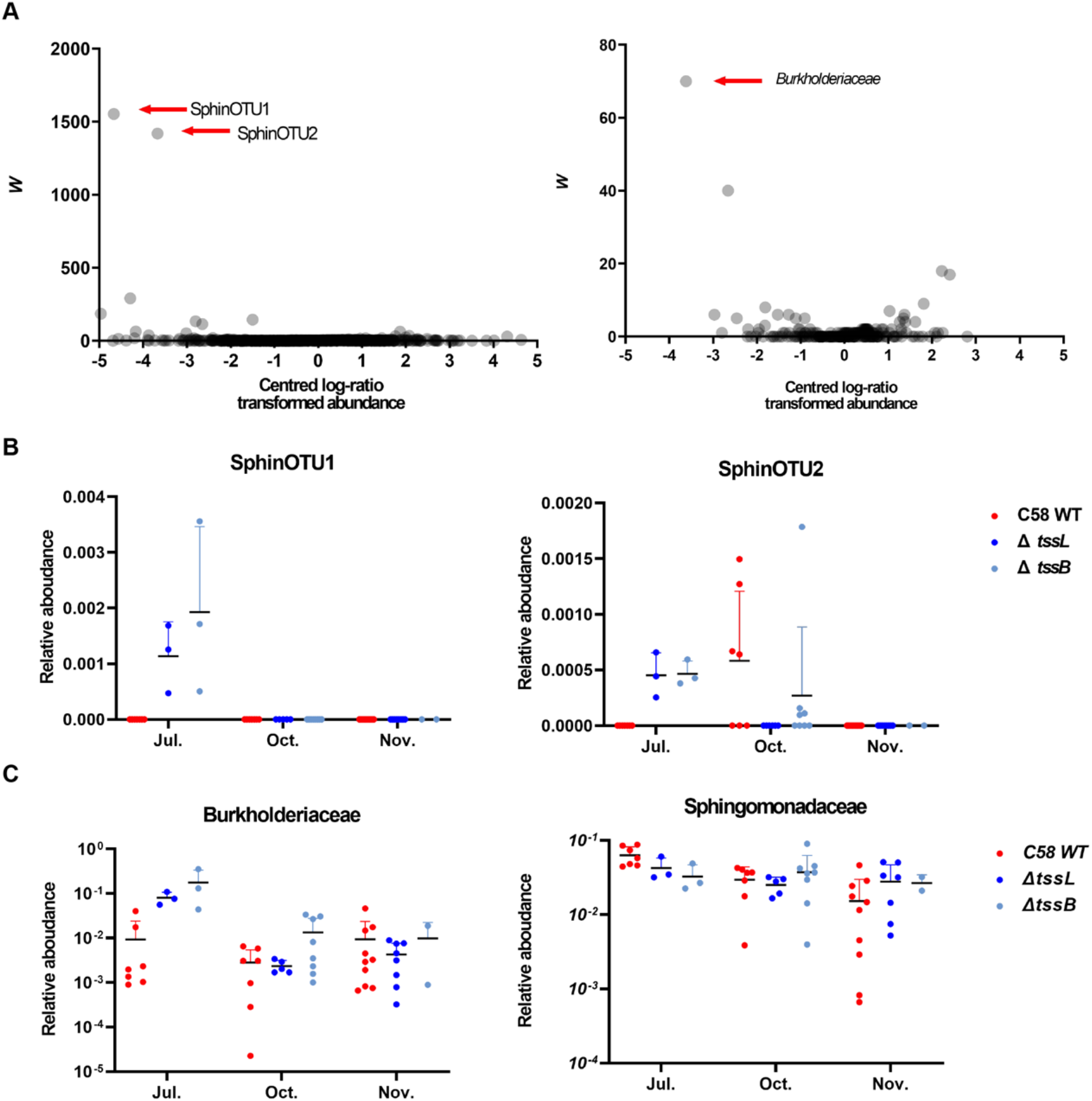
Analysis of composition of microbiomes (ANCOM) for gallobiomes in July. The ANCOM volcano plots were drawn based on the centred-log-ratio-transformed abundance of (A) OTUs or bacterial families. The taxa that exhibit differential abundance (DA) between gallobiomes induced by C58 WT and T6SS mutants are indicated by red arrows. The two DA OTUs belonging to *Sphingomonadaceae* were named SphinOTU1 and SphinOTU2, respectively. The relative abundance of focal taxa was shown in panels (B) and (C). Lines and error bars indicate mean ± SD.

### T6SS-dependent antagonism between agrobacteria and a *Sphingomonas* sp. isolate

The T6SS-dependent differential abundance of two *Sphingomonas* OTUs in gallobiomes motivated us to investigate whether C58 exhibits T6SS antibacterial activity to *Sphingomonas*. A *Sphingomonas* sp. strain R1 isolated from the tomato rhizosphere in one soil inoculation experiment was used for interbacterial competition assays *in planta* and *in vitro*. Each of the agrobacterial strains was mixed at 1:1 ratio with *Sphingomonas* sp. R1 as inoculum and perform soil inoculation on wounded tomato seedlings. Colonization efficiency of those strains was determined by counting colony forming unit (CFU) recovered from wounded stem segments at 10 day-post-inoculation (dpi). The results showed a ~ 0.5 log of reduced CFUs of *Sphingomonas* sp. R1 when co-inoculated with WT as compared to that with Δ*tssL* and Δ*tssB* or R1 only (Fig. 7A). Recovered CFUs of agrobacterial WT and T6SS mutants inoculated alone or with *Sphingomonas* sp. R1. are 1 to 1.5 log higher than *Sphingomonas* sp. R1 but no difference among agrobacterial WT and T6SS mutants inoculated alone or with *Sphingomonas* sp. R1. These results suggest that agrobacteria have higher competitive colonization efficiency than *Sphingomonas* sp. R1 in part dependent on a functional T6SS. Accordingly, interbacterial competition assay on agar plate also showed T6SS-dependent antibacterial activity to *Sphingomonas* sp. R1 (Fig. 7B). However, when interbacterial competition was carried out *in vitro* on agar plates, only weak antibacterial activity was observed at 1:1 ratio but ~0.5 log reduced CFUs of *Sphingomonas* sp. R1 was detected when competition was carried out at 10:1 ratio of agrobacteria: *Sphingomonas* sp. R1. The results show that the T6SS provides agrobacteria with a higher competitive advantage against *Sphingomonas* sp. R1 on plant wounding site than *in vitro*. We further evaluated whether CFUs of *Sphingomonas* sp. R1 was enriched in crown galls induced by T6SS mutants as compared to those induced by WT via soil inoculation method used for 16S rRNA gene amplicon sequencing. Surprisingly, *Sphingomonas* sp. R1 was not always present in crown galls at 28 dpi, in which R1 was only recovered from one out of three independent experiments by direct induction of crown galls on tomato stem (Fig. S3). The abundance of C58 ranged from 10^4^ to 10^7^ CFU per gall and no consistent difference could be observed between the WT and the T6SS mutants. Together, these results suggest that C58 exhibits antibacterial activity to inhibit the *in vitro* growth and plant colonization of *Sphingomonas* sp. R1. The T6SS may help agrobacteria to gain competitive growth over other competing rhizobacteria on plant surface for effective infection, leading to higher disease incidence.

**Fig. 7.**
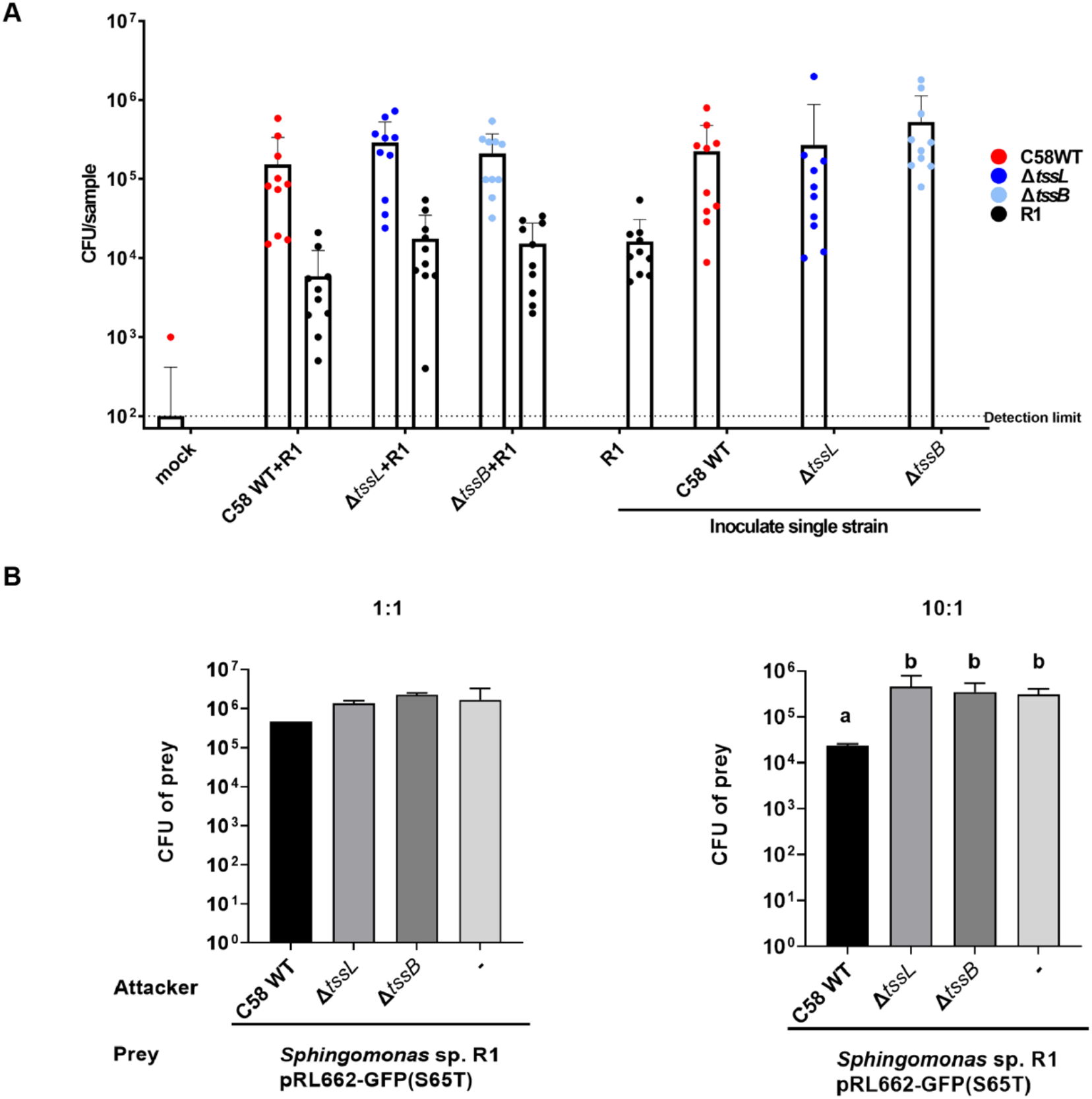
Competition between C58 and *Sphingomonas* sp. R1 *in vitro* and on wounded site of tomato stems. Each of the three *Agrobacterium* strains (i.e., C58 WT, Δ*tssL*, and Δ*tssB*) was individually tested for competition against *Sphingomonas* sp. R1 isolated from rhizosphere of tomato. (A) *Agrobacterium* -only or co-infected with *Sphingomonas* sp. R1 at 1:1 ratio in soil were recovered from the surface of wounded stem segments at 10 dpi, the CFU of *Agrobacterium* and *Sphingomonas* sp. R1 were plotted. T-test indicated the CFU differences of *Sphingomonas* sp. R1 inoculation alone or co-infection with T6SS mutants as compared to WT with indicated *p* values. Data are mean ± SD of three independent experiments, each with five seedlings inoculated for each strain. ANOVA test indicated no significant difference among means (*p*-value = 0.0945 and 0.0551 for *Agrobacterium* and *Sphingomonas* sp. R1 CFUs, respectively). (B) The *in vitro* competition results based on *Agrobacterium*:*Sphingomonas* sp. R1 at 1:1 and 10:1 ratio, and the survival of *Sphingomonas* sp. R1 was plotted; lines and error bars indicate mean ± SD of four biological replicates from two independent experiments. Based on ANOVA followed by Tukey’s multiple comparison test, C58 WT significantly reduced *Sphingomonas* sp. R1 CFUs when mixed in 10:1 ratio (*p*-value = 0.0153) but not in 1:1 ratio (*p*-value = 0.2753).

## DISCUSSION

In this study, we designed experiments to address whether the T6SS affects agrobacterial tumorigenesis and gallobiome composition. By establishing a soil inoculation platform to evaluate the disease incidence of wounded tomato seedlings across seasons, we revealed that a functional T6SS is positively correlated with disease occurrence but not gall weight. Furthermore, by developing an effective protocol for enrichment of bacterial 16S rRNA genes by utilizing blocker to prevent host contamination, we found that seasons or environmental factors are the major driver in shaping gallobiome composition while T6SS could also influence microbiota at more specific manner.

The evidences that agrobacterial T6SS promotes tumorigenesis in soil inoculation (Table 1, Fig. 1) but not direct inoculation (19) suggest that the T6SS is not directly involved in virulence. Instead, agrobacteria may deploy the antibacterial weapon to increase the occurrence of infection in the presence of relatively complex microbial communities in rhizosphere (Table S3, (26)). Once galls are induced, the T6SS is not required for gall development since no significant difference in weight was observed between WT and T6SS mutants (Fig. 1B). The presence of a functional T6SS has no impact on the population size of agrobacteria, which is consistent with recent findings that T6SS genes were not identified as fitness genes by Tn-seq screen in crown galls (27). The T6SS of C58 also appears not to be the key factor to shape gallobiome composition, suggesting that agrobacteria may also use other ways for niche competition in galls. It is generally believed that the agrobacteria inciting crown galls and later become residents have privilege to access the specific opines synthesized by transformed plant cells containing the agrobacterial T-DNA (28). This “opine concept” has not been experimentally validated but supported by a study showed that the ability of agrobacteria to trap opines can acquire a competitive advantage over the siblings which cannot utilize opine (29). Therefore, it would be interesting to investigate whether opine synthesis and other fitness genes would impact the crown gall microbiota in which T6SS may synergistically enhance the niche occupation.

The findings that disease incidence and crown gall weight are reversely correlated with temperature (Fig. 1C) are expected. Early studies demonstrated that crown gall sizes were dramatically decreased when the host plants were inoculated at 28-30°C, while no gall formation at 31°C or above (30, 31). In addition, the T4SS complex and its associated T-pilus are more stable or produced at higher levels at 19°C than 28°C whereas T4SS-mediated plasmid conjugation is deficient at 28°C (32–36). The increased tumor weight when inoculated in fall/winter is also correlated with a higher relative abundance of agrobacterial OTU (Fig. 5G), suggesting the active proliferation of agrobacterial population inside crown galls. Furthermore, our data show that crown galls produced in summer have distinct microbiota as compared to those from fall/winter (Fig. 4B), which echoes previous studies on *Allorhizobium vitis* that crown gall microbiota of grapevine is significantly different in summer (37). Interestingly, while there is no global difference of gallobiome induced by WT or T6SS mutants inoculated in fall/winter, we found that the two *Sphingomonas* OTUs and the family Burkholderiaceae were only present or significantly enriched in the gallobiome induced by T6SS mutants in summer (Fig. 6). By counting viable agrobacterial inoculum, we showed that C58 exhibits a T6SS-mediated antibacterial activity to *Sphingomonas* sp. R1 *in vitro* and in rhizosphere (Fig. 7A, B) but no significant difference of R1 abundance between the galls co-infected with C58 WT or T6SS mutants (Fig. S3). Interestingly, we noticed that the colonization efficiency of those T6SS mutants was more variable as compared to those of WT from the combined data set (Fig. S4). To this end, it remains to be determined whether the absence or reduced abundance of *Sphingomonas* and Burkholderiaceae of gallobiome induced by WT as compared to those by T6SS mutants are direct consequence of antibacterial activity conferred by T6SS. However, current evidence suggests that agrobacteria may use T6SS to gain competitive advantage by its antibacterial activity against other bacterial species residing in the soil or rhizosphere and therefore increases its success in inciting tumors on infected plants.

Besides Rhizobiaceae that agrobacteria belong to, other bacterial families were also occupied considerable proportions of crown gall microbiota, such as Comamonadaceae, members of which are commonly found in soil and water as well as plant-associated bacteria *Variovorax* shown to be important for root growth of *Arabidopsis* (38, 39). Bacteria belong to Xanthobacteraceae and Pseudomonaceae, the two families highly associated with plants as either pathogens or commensals, also had high relative abundance in the crown galls. Some *Pseudomonas* species such as *P. fluorescens* and *P. putida* are capable of utilizing opines (40, 41), which may explain why bacterial species other than the agrobacterial inoculum could also colonize and propagate in high relative abundance in crown galls.

We found that the predominance of plant DNA in crown gall samples led to the low resolution of endophytic bacterial microbiota survey via standard 16S rRNA gene amplicon sequencing due to the co-amplification of chloroplast and mitochondrial sequences. Although the use of 16S rRNA gene primers containing mismatches to 16S rRNA gene of plant chloroplast or mitochondria could potentially reduce host contamination in endophytic microbiota studies (25, 26), the “bacteria-specific” primers resulted in > 90% host reads of our initial attempt (Round I) (Table 2), which is similar to results of a previous study (22). When adding blockers that could specifically anneal to chloroplast or mitochondrial DNA but not be extended by DNA polymerase during PCR, we successfully reduced the host contamination and obtained sufficient sequencing depth for this study. Our results support the effectiveness of blockers in increasing bacterial reads of endophyte community (22–24). Our newly developed SI-BBacSeq protocol for gallobiome may be applicable to microbiota associated with tomato, or even other plants after modifying the blocker sequences.

To date, collective plant microbiota studies suggest the enrichment of T6SS genes in microbiota and as a trait for niche competition in microbiota (5, 8). However, the role and impact of T6SS in different plant and microbiome contexts may be different. Our study suggests that the T6SS may provide agrobacteria with competitive advantages at initial stage of infection but likely not critical for proliferation once they established their niche inside the crown galls. Future works that investigate the spatial distribution of agrobacterial cells and other endophytes inside crown galls or plant wounding site, together with temporal and spatial visualization of the T6SS activity, are important for dissecting the roles of T6SS in agrobacterial disease ecology.

## MATERIALS AND METHODS

### Bacterial strains and growth condition

Bacterial strains and plasmids used in this study are listed in Table S4. *Agrobacterium* strain C58 wild type and two T6SS mutants (i.e., Δ*tssB* and Δ*tssL*) (42) were first streaked on 523 medium agar plates (43) and incubate at 25 °C for 48 hr. *Sphingomonas* sp. R1 was grown on R2A medium without soluble starch and pyruvate (44). *Escherichia coli* was grown on lysogeny broth (LB) medium at 37°C. Freshly grown colony was inoculated into corresponding broth for overnight culture. The antibiotics and concentration used are: 10 μg/ml gentamycin for *E. coli*, 50 μg/ml gentamycin for *Agrobacterium*, and 20 μg/ml kanamycin and 50μg/ml gentamycin for *Sphingomonas* sp. R1.

### Construction of strains enable antibiotics selection

Primers used for construction are listed in Table S5. The DNA fragment containing gentamycin resistance (Gm^R^) gene and green fluorescence protein *gfp*(S65T) were amplified from pRL662::GFP(S65T) with the primer pair “BclI-GFP-GmR-F-25” and “BclI-GFP-GmR-R-25”, and then cloned into XbaI digested pJQ-COM. The newly constructed plasmid was named pJQ-com-GmR-GFP, which enabled the generation of Gm^R^ gene and *gfp* (S65T) knocked in at the upstream of *actC* (*atu5330*) of *Agrobacterium* genome after double crossover. The knock-in strains were generated in *Agrobacterium* C58, Δ*tssB*, and Δ*tssL* background. The knock-in event and excision of pJQ200ks backbone were confirmed by GFP signal, gentamycin resistance, and PCR with the primer pair “3’sacB-5’GmR pJQ200 F” and “3’sacB-5’GmR pJQ200 R”, which target the fragment between *sacB* and Gm^R^ gene on pJQ200ks. The plasmids pRL66::GFP(S65T) and pBBR::mCherry conferring kanamycin resistance were each transformed into *Sphingomonas* sp. R1 through electroporation. The procedure of electroporation, doble crossover via pJQ200ks were performed as described previously (45).

### Plant material and growth condition

Seeds of *Solanum lycopersicum* (tomato) cultivar Known-You 301 from KNOWN-YOU SEED CO., LTD (Kaohsiung, Taiwan) were germinated and grown in unsterilized potting mixture (Jiffy^®^Premium Fine Peat Substrate, perlite, and vermiculite, mixed in 4:1:1 ratio, v:v:v) in the greenhouse EL329 (N25.043047890856812, E121.61135464167913, Greenhouse building, Academia Sinica, Taipei, Taiwan).

### Soil inoculation

The soil inoculation method was optimized based on the method used for agrobacterial infection of pea and walnut (46, 47). Overnight (14-16 hr) culture of agrobacterial strain in 12 ml of 523 broth were centrifugated at 6,000 × g for 10 min. The pellets were washed once by using 10 ml of sterile saline (0.9% NaCl in H_2_O). The washed pellets were centrifuged and resuspended in sterile saline with OD_600_ adjusted to 1. Bacterial suspension was mixed into unsterilized soil (1 ml suspension per 100 g of soil), which is expected to result in ~10^7^ colony-forming units (CFUs)/g soil, the optimized concentration that could incite crown galls but not at 100%. Tomato seedlings with two true leaves (two-to three-week-old) were wounded at the site between primary root and cotyledons by a fire-sterilized sewing needle, planted into the inoculated soil, and grown in the greenhouse. Each strain was inoculated with 14 to 20 tomato seedlings, and the inoculation with saline were used as a negative control.

### Harvest, surface sterilization and storage of crown gall samples

Two months after the inoculation, tomato plants were harvested to examine gall formation on the wounded site. The disease incidence was calculated by dividing the number of plants with visible gall formation by the total number of inoculated plants. The sections with crown gall were cut out and sterilized in 35 ml of sterilization solution (3% NaOCl, 0.01% Tween-20) for 30s, then transferred to 35 ml of 70% ethanol and washed in 35 ml of sterile H_2_O for three times (48). The sterilization was evaluated by spreading 100 μl liquid from the third wash onto 523 agar plate for 2 days to make sure that there was no growth of bacteria. After sterilization, the crown galls were dissected from each of the plant segments in sterile petri dish, wrapped in sterile aluminum foil, frozen by liquid nitrogen, and stored at −80 °C prior to DNA extraction.

### DNA extraction form crown galls

Crown gall samples were homogenized by pestle and mortar with liquid nitrogen. 0.25 g of homogenized tissue were transferred to the beating tube of DNeasy^®^PowerSoil^®^Kit (Qiagen, Germany) following the manufacturer’s instructions. Concentrations of extracted DNA were determined by NanoDrop^™^ 1000 (Thermo Fisher Scientific, United State) and Qubit dsDNA HS (Invitrogen, United State).

### 16S rRNA gene amplification and sequencing

Three rounds of amplicon sequencing were conducted in this study. The round I was conducted according to established protocols of soil microbiota studies using 16S rRNA gene variable regions (25). Each primer with a partial adaptor sequence was compatible to Illumina Truseq Combinatorial Dual index system. The first PCR amplifications were carried out in following 25 μl reaction: 5 ng DNA template, 2× KAPA HiFi HotStart DNA Polymerase ReadyMix (Roche, Switzerland), and 0.2 μM for each of the forward and reverse primers. The first PCR program and reaction details for V3-4 and V5-7 are listed in Table S6. The PCR products were purified by Ampure XP beads (Thermo Fisher Scientific Inc, Sweden). The V5-7 PCR products underwent BluePippin (Sage Science) size-selection to remove tomato amplicon. To attach dual indices and full-length Illumina sequencing adapters, a second round of PCR amplifications were employed for 50 μl reactions with purified or size-selection PCR products, 2× KAPA HiFi HotStart DNA Polymerase ReadyMix (Roche, Switzerland), and 0.5 μM for each of the forward and reverse primers. All the library products of second PCR were purified and pooled. Pooled libraries were loaded onto an Illumina MiSeq V3 flow cell (Illumina, United States) and for 2×300 bp paired-end sequencing.

The round II of amplicon sequencing was conducted to test the effect of adding PCR blockers on reducing plant chloroplast and mitochondrial sequences. Three gall samples (1115-W21, 1115-W22 and 1115-W25) were used as the DNA templates to provide biological replicates. For each DNA sample, four primer pairs targeting different variable regions of 16S rRNA gene were used, including: V1-3, V3-4, V5-7, and V6-8 (Table S1). Each primer contained a partial adaptor sequence of the Illumina Truseq Combinatorial Dual index system. Cognate 3’ C3 spacer modified oligonucleotides (blockers) which would block the amplification of tomato chloroplast and mitochondrial 16S rRNA genes were designed according to a previous study (22) and applied to one of the two PCRs for each sample-primer combination (Table S2). Procedures for PCR and Illumina sequencing were based on those used for round I, but the amount of DNA template was increased to 25 ng to reduce PCR cycle time and the annealing temperature was adjusted for blockers. More detailed technical information is provided in Table S6. Based on results from the first two rounds of amplicon sequencing, an optimized protocol that utilizes blockers and targets V5-7 regions was used for round III. All 53 crown gall samples were included in this final round.

### Raw read processing and microbiota analysis

The Illumina raw reads in FASTQ format were imported into QIIME2 release 2020.8 (49). The primer region of imported sequences was trimmed using DADA2 (50). The denoised and merged paired-end reads were used to construct output feature tables containing amplicon sequence variants (ASVs) ID and counts in different samples. For amplicon sequencing round III, the ASVs were further clustered into species-level OTUs based on 99% sequence identity using the “qiime vsearch cluster-features-de-novo” function (51) and the singletons were removed by using the “qiime feature-table filter-features” function prior to the downstream analysis. The taxonomic information was inferred for representative OTU sequences via pre-trained q2-feature-classifier (52) and classify-sklearn naïve Bayes classifier based on SILVA version 138 SSU Ref NR 99 full-length sequences dataset (53). The sequences assigned as chloroplasts, mitochondria, eukaryotes, archaea, or unknown were removed. After the filtering, a phylogenetic tree of eubacterial sequences was inferred using the q2-fragment-insertion plug-in (54–57) based on the SILVA 128 SEPP reference database. Weighted Unifrac (58) analysis of those datasets was performed based on insertion tree; the distance matrix and principal coordinate analysis (PCoA) plot were generated using QIIME2. For round III, phylogenic and diversity analysis was conducted using “qiime diversity core-metrics-phylogenetic” and the sub-sampling depth was set to 9,800 reads/sample.

### Differential abundance analysis of OTU

Analysis of composition of microbiomes (ANCOM) was applied to identify OTUs that have significantly different relative abundance in the gallobiome associated with C58 WT and T6SS mutants using the function “qiime composition ancom” in QIIME2. The feature table collapsed into family level was imported and transformed by centred log-ratio transformation. The null hypothesis of ANCOM is that the abundance of a OTU is not different between two study groups. After performing all comparison between each OTU in two study groups, the times that the null hypothesis was rejected is called *w*. The OTUs having *w* values at 70 percentile or higher are considered as significant (59, 60).

### Isolation of *Sphingomonas* sp. R1

*Sphingomonas* sp. R1 was isolated from a wounded tomato stem during a soil inoculation assay. After 10 dpi of soil inoculation mentioned above, the 1-cm wounded segment of tomato seedlings were harvested and rinsed by sterile water, and then transferred to 1 ml 0.9% NaCl. After vortex at max speed for 3 minutes, the supernatant was transferred to a new microcentrifuge tube and spread on 523 agar plates via exponential mode of easySpiral Dilute^®^ (Ref. 414 000, INTERSCIENCE, France). After growth at 25°C for two days, ~ 20 colonies were selected and streaked out on 523 agar plates for pure culture. The partial 16S rRNA gene of the isolates was amplified and sequenced by V5-V7 primer set, which was used to BLAST in 16S ribosomal RNA gene sequence database in NCBI for identification of 16 bacterial species (Table S3).

### Interbacterial competition assay *in vitro* and *in planta*

For interbacterial competition *in vitro*, the bacterial suspension was adjusted to OD_600_ equals to 3.0 in 0.9% NaCl (w/v) after overnight culture. For the *in vitro* competition, the attacker strains (C58 WT, Δ*tssL*, or Δ*tssB*) were further diluted into OD_600_ equals to 1, and the prey *Sphingomonas* sp. R1 containing pRL662::GFP was further diluted into OD_600_ equal to 0.3 or 0.1 for mixing with the attacker in equal volume for different density ratio. Two spots of 10 μl mixture were spotted on *Agrobacterium* kill-triggering medium (21). After co-incubated for 16 h in 25°C, the colonies were scraped and resuspended in 1 ml 0.9% NaCl (w/v), and spread via exponential mode of easySpiral Dilute^®^ (Ref. 414 000, INTERSCIENCE, France) on 523 media containing 30 μg/ml gentamycin to recover the prey.

For agrobacterial colonization with or without *Sphingomonas* sp. R1, each of the agrobacterial strains (C58 WT, Δ*tssL* or Δ*tssB*) with Gm^R^-GFP knock-in was mixed with *Sphingomonas* sp. R1 containing pBBR::mCherry at equal OD_600_ =1.0. The 5 μl mixture was inoculated on the needle-wounded stem of tomato seedling with first pair of true leaves. Tomato seedlings were harvested at 10 days post-inoculation (dpi). For each plant, the 1 cm wounded segment was cut and rinsed by sterile water before being transferred into a sterilized Eppendorf tube containing 1 ml of 0.9% NaCl. After vortex at max speed for 3 minutes, the liquid was considered as the rhizosphere sample. The samples were diluted and plated on media with gentamycin for CFU calculation.

When counting the CFU of C58 and *Sphingomonas* sp. R1 in the tumor, the crown galls were collected after growing in the greenhouse for three weeks. The 1-cm tomato stem section containing the gall in middle was cut and homogenized with 1 ml 0.9% NaCl (w/v) by sterilized mortar and pestle. The homogenized tissue was diluted 10-fold and spread via exponential mode of easySpiral Dilute^®^ on 523 media containing 50 ppm gentamycin to recover the attacker and prey. After incubation for two days at 25°C, the colonies were counted by using the Scan^®^ 500 Automatic colony counter (Ref 436 000, INTERSCIENCE, France; software version 8.6.1) to calculate CFUs.

### Statistics and visualization

GraphPad Prism 8 was used to perform t-test on sequencing read counts, one-way ANOVA on CFU counts, two-way ANOVA followed by Turkey’s multiple comparison test on crown gall disease incidence and alpha diversity index, Kruskai-Wallis test on tumor weight, Spearman correlation on gall weight and relative abundance of agrobacterial OTU. The alpha-rarefaction curves were plotted using Phyloseq (61) in R version 3.6.2. ADONIS in “qiime diversity beta-group-significance” of QIIME2 (62, 63) was used to test if groups of samples have significantly different microbiota composition. The taxonomic composition plots were generated using qiime2R (v0.99.35, https://github.com/jbisanz/qiime2R) in R studio 2.

## DATA AVAILABILITY

All raw data sets are available in the National Center for Biotechnology Information (NCBI) under BioProject accession PRJNA894311.

## AUTHOR CONTRIBUTION

Conceptualization: SCW, CHK, EML

Funding acquisition: CHK, EML

Investigation: SCW, CHK, EML Methodology: SCW, APC, SJC, CHK

Project administration: EML

Supervision: CHK, EML

Validation: SCW, CHK, EML

Visualization: SCW, CHK, EML

Writing – original draft: SCW, CHK, EML

Writing – review & editing: SCW, APC, SJC, CHK, EML

## ACKNOWLEDGEMENTS

The authors thank Teng-Kuei Huang and Mao-Sen Liu for careful reading of this manuscript and the members of Lai and Kuo labs for helpful discussion. We also acknowledge Lay-Sun Ma for constructing pRL662::GFP(S65T) and Stanton Gelvin for providing pBBR::mCherry. The Illumina sequencing library preparation was carried out in the Genomic Technology Core (Institute of Plant and Microbial Biology, Academia Sinica). We thank the Illumina MiSeq sequencing service provided by the Genomics Core (Institute of Molecular Biology, Academia Sinica).

## FIGURE LEGENDS

**Figure S1. DNA sequence alignment of 16S rRNA genes of selected bacterial strains and tomato KnowYou 301 chloroplast and mitochondria.**

The 16S rRNA gene sequences of chloroplasts and mitochondria in *Solanum lycopersicum* (tomato) cultivar Known-You 301 were obtained from this study. All of the bacterial 16S rRNA gene sequences were accessed from Reference Sequence (RefSeq) database in National Center of Biotechnology Information (NCBI). The species, strain name, and accession number of 16S rRNA gene were listed on the left of each sequence. The alignment of sequences mentioned above were aligned in MEGAX via ClustalW multiple alignment with gap opening penalty 15.00 and gap extension penalty 6.66 (default). The levels of shaded blue color reflect the degree of identity. The regions of each primer and blocker were underlined. The sequences highlighted by red, black, yellow and green are the annealing region of primer sets for V1-3, V3-4, V5-7 and V6-8 in 16S rRNA gene, respectively. And the gray framed sequences in tomato chloroplast and mitochondrial 16S rRNA genes are the blockers for cognate primers.

**Figure S2. Alpha rarefaction curves of the observed bacterial OTUs based on amplicon sequencing of 53 gall samples in sequencing run III.**

The rarefaction curves were plotted after clustering DADA2-output ASVs into 99% OTUs. Sample size in X-axis indicates different sub-sampling depth of each dataset; Y-axis indicates the number of observed OTUs under certain sub-sampling depth. Label of each curves indicates the sample ID in metadata.

**Figure S3.Competition of** *Agrobacterium* **and *Sphingomonas* sp. R1 in tomato gall on stem at 28 dpi**.

Each of the three *Agrobacterium* strains (i.e., C58 WT, Δ*tssL*, and Δ*tssB*) and R1 were mixed at 1:1 ratio then inoculated on wounded tomato stem. The galls were harvested and homogenized for plating at 28 dpi. *Agrobacterium* strains and *Sphingomonas* sp. R1 were recovered on 523 medium plates containing proper antibiotics. CFU data for agrobacteria are mean ± SD of three independent experiments/batches, each with five seedlings inoculated for each strain. R1 could only be recovered from galls in batch 3. The *p*-value of ANOVA against CFU numbers of C58 for each batch is 0.786, 0.0023 and 0.230; the *p*-value of ANOVA against CFU numbers of R1 was 0.6413. The significance of CFU numbers of C58 in second batch is due to the low CFU counts of Δ*tssL* single inoculation group, which could not be replicated in other two batches.

**Figure S4. Agrobacteria colonization on wounded stem segments.**

The colonies of *Agrobacterium* strains C58 WT, Δ*tssL*, and Δ*tssB* recovered from the surface of wounded stem segments following the soil inoculation procedure were plotted. Different symbols indicated the outcome from different batches of colonization assay. *F*-test indicated the variance of Δ*tssL* and Δ*tssB* were significantly different when compared to C58 WT (*p*(*F*≤*f*) = 0.00028 and 1.77826E-05, respectively). Brown-Forsythe and Welch ANOVA test indicated no significant difference among means (*p-*value= 0.4847). Line indicates median.

## REFERENCES

1. Coulthurst S. 2019. The Type VI secretion system: a versatile bacterial weapon. Microbiology (Reading) 165:503–515.

2. Jurenas D, Journet L. 2021. Activity, delivery, and diversity of Type VI secretion effectors. Mol Microbiol 115:383–394.

3. Lien YW, Lai EM. 2017. Type VI Secretion Effectors: Methodologies and Biology. Front Cell Infect Microbiol 7:254.

4. Gallegos-Monterrosa R, Coulthurst SJ. 2021. The ecological impact of a bacterial weapon: microbial interactions and the Type VI secretion system. FEMS Microbiol Rev 45.

5. Bulgarelli D, Garrido-Oter R, Munch PC, Weiman A, Droge J, Pan Y, McHardy AC, Schulze-Lefert P. 2015. Structure and function of the bacterial root microbiota in wild and domesticated barley. Cell Host Microbe 17:392–403.

6. Wexler AG, Bao Y, Whitney JC, Bobay LM, Xavier JB, Schofield WB, Barry NA, Russell AB, Tran BQ, Goo YA, Goodlett DR, Ochman H, Mougous JD, Goodman AL. 2016. Human symbionts inject and neutralize antibacterial toxins to persist in the gut. Proc Natl Acad Sci U S A 113:3639–44.

7. Verster AJ, Ross BD, Radey MC, Bao Y, Goodman AL, Mougous JD, Borenstein E. 2017. The Landscape of Type VI Secretion across Human Gut Microbiomes Reveals Its Role in Community Composition. Cell Host Microbe 22:411–419 e4.

8. Vogel CM, Potthoff DB, Schafer M, Barandun N, Vorholt JA. 2021. Protective role of the Arabidopsis leaf microbiota against a bacterial pathogen. Nat Microbiol 6:1537–1548.

9. Serapio-Palacios A, Woodward SE, Vogt SL, Deng W, Creus-Cuadros A, Huus KE, Cirstea M, Gerrie M, Barcik W, Yu H, Finlay BB. 2022. Type VI secretion systems of pathogenic and commensal bacteria mediate niche occupancy in the gut. Cell Rep 39:110731.

10. Ma LS, Hachani A, Lin JS, Filloux A, Lai EM. 2014. Agrobacterium tumefaciens deploys a superfamily of type VI secretion DNase effectors as weapons for interbacterial competition in planta. Cell Host Microbe 16:94–104.

11. Bernal P, Allsopp LP, Filloux A, Llamas MA. 2017. The Pseudomonas putida T6SS is a plant warden against phytopathogens. ISME J 11:972–987.

12. Bernal P, Llamas MA, Filloux A. 2018. Type VI secretion systems in plant-associated bacteria. Environ Microbiol 20:1–15.

13. Vacheron J, Pechy-Tarr M, Brochet S, Heiman CM, Stojiljkovic M, Maurhofer M, Keel C. 2019. T6SS contributes to gut microbiome invasion and killing of an herbivorous pest insect by plant-beneficial Pseudomonas protegens. ISME J 13:1318–1329.

14. de Lajudie PM, Andrews M, Ardley J, Eardly B, Jumas-Bilak E, Kuzmanovic N, Lassalle F, Lindstrom K, Mhamdi R, Martinez-Romero E, Moulin L, Mousavi SA, Nesme X, Peix A, Pulawska J, Steenkamp E, Stepkowski T, Tian CF, Vinuesa P, Wei G, Willems A, Zilli J, Young P. 2019. Minimal standards for the description of new genera and species of rhizobia and agrobacteria. Int J Syst Evol Microbiol 69:1852–1863.

15. Hwang HH, Yu M, Lai EM. 2017. Agrobacterium-Mediated Plant Transformation: Biology and Applications, vol 15. American Society of Plant Biologists.

16. Wu CF, Santos MNM, Cho ST, Chang HH, Tsai YM, Smith DA, Kuo CH, Chang JH, Lai EM. 2019. Plant-Pathogenic Agrobacterium tumefaciens Strains Have Diverse Type VI Effector-Immunity Pairs and Vary in In-Planta Competitiveness. Molecular Plant-Microbe Interactions 32:961–971.

17. Wu CF, Weisberg AJ, Davis EW, 2nd, Chou L, Khan S, Lai EM, Kuo CH, Chang JH. 2021. Diversification of the Type VI Secretion System in Agrobacteria. mBio 12:e0192721.

18. Chou L, Lin YC, Haryono M, Santos MNM, Cho ST, Weisberg AJ, Wu CF, Chang JH, Lai EM, Kuo CH. 2022. Modular evolution of secretion systems and virulence plasmids in a bacterial species complex. BMC Biol 20:16.

19. Wu HY, Chung PC, Shih HW, Wen SR, Lai EM. 2008. Secretome Analysis Uncovers an Hcp-Family Protein Secreted via a Type VI Secretion System in *Agrobacterium tumefaciens*. J Bacteriol 190:2841–2850.

20. Lassalle F, Campillo T, Vial L, Baude J, Costechareyre D, Chapulliot D, Shams M, Abrouk D, Lavire C, Oger-Desfeux C, Hommais F, Gueguen L, Daubin V, Muller D, Nesme X. 2011. Genomic species are ecological species as revealed by comparative genomics in Agrobacterium tumefaciens. Genome Biol Evol 3:762–81.

21. Yu M, Wang YC, Huang CJ, Ma LS, Lai EM. 2020. Agrobacterium tumefaciens Deploys a Versatile Antibacterial Strategy to Increase its Competitiveness. J Bacteriol doi:10.1128/JB.00490-20.

22. Lefevre E, Gardner CM, Gunsch CK. 2020. A novel PCR-clamping assay reducing plant host DNA amplification significantly improves prokaryotic endo-microbiome community characterization. Fems Microbiology Ecology 96.

23. Fitzpatrick CR, Lu-Irving P, Copeland J, Guttman DS, Wang PW, Baltrus DA, Dlugosch KM, Johnson MTJ. 2018. Chloroplast sequence variation and the efficacy of peptide nucleic acids for blocking host amplification in plant microbiome studies. Microbiome 6:144.

24. Arenz BE, Schlatter DC, Bradeen JM, Kinkel LL. 2015. Blocking primers reduce co-amplification of plant DNA when studying bacterial endophyte communities. Journal of Microbiological Methods 117:1–3.

25. Thijs S, Op De Beeck M, Beckers B, Truyens S, Stevens V, Van Hamme JD, Weyens N, Vangronsveld J. 2017. Comparative Evaluation of Four Bacteria-Specific Primer Pairs for 16S rRNA Gene Surveys. Front Microbiol 8:494.

26. Beckers B, Op De Beeck M, Thijs S, Truyens S, Weyens N, Boerjan W, Vangronsveld J. 2016. Performance of 16s rDNA Primer Pairs in the Study of Rhizosphere and Endosphere Bacterial Microbiomes in Metabarcoding Studies. Front Microbiol 7:650.

27. Torres M, Jiquel A, Jeanne E, Naquin D, Dessaux Y, Faure D. 2022. Agrobacterium tumefaciens fitness genes involved in the colonization of plant tumors and roots. New Phytol 233:905–918.

28. Guyon P, Chilton MD, Petit A, Tempe J. 1980. Agropine in “null-type” crown gall tumors: Evidence for generality of the opine concept. Proc Natl Acad Sci U S A 77:2693–7.

29. Lang J, Vigouroux A, Planamente S, El Sahili A, Blin P, Aumont-Nicaise M, Dessaux Y, Morera S, Faure D. 2014. Agrobacterium uses a unique ligand-binding mode for trapping opines and acquiring a competitive advantage in the niche construction on plant host. PLoS Pathog 10:e1004444.

30. Riker AJ. 1926. Studies on the influence of some environmental factors on the development of crown gall. Journal of Agricultural Research 32:0083–0096.

31. Braun AC. 1947. Thermal Studies on the Factors Responsible for Tumor Initiation in Crown Gall. American Journal of Botany 34:234–240.

32. Fullner KJ, Nester EW. 1996. Temperature affects the T-DNA transfer machinery of Agrobacterium tumefaciens. J Bacteriol 178:1498–504.

33. Fullner KJ, Lara JC, Nester EW. 1996. Pilus assembly by Agrobacterium T-DNA transfer genes. Science 273:1107–9.

34. Lai EM, Kado CI. 1998. Processed VirB2 is the major subunit of the promiscuous pilus of Agrobacterium tumefaciens. J Bacteriol 180:2711–7.

35. Lai EM, Kado CI. 2000. The T-pilus of Agrobacterium tumefaciens. Trends Microbiol 8:361–9.

36. Baron C, Domke N, Beinhofer M, Hapfelmeier S. 2001. Elevated temperature differentially affects virulence, VirB protein accumulation, and T-pilus formation in different Agrobacterium tumefaciens and Agrobacterium vitis strains. J Bacteriol 183:6852–61.

37. Faist H, Keller A, Hentschel U, Deeken R. 2016. Grapevine (Vitis vinifera) Crown Galls Host Distinct Microbiota. Appl Environ Microbiol 82:5542–52.

38. Willems A, De Ley., J., Gillis, M. Kersters, D. K.. 1991. Comamonadaceae, a New Family Encompassing the Acidovorans rRNA Complex, Including Variovoraxparadoxus gen. nov., comb. nov. for Alcaligenes paradoxus (Davis 1969). INTERNATIONJOAULRNAOLF SYSTEMATBICACTERIOLO 41.

39. Finkel OM, Salas-Gonzalez I, Castrillo G, Conway JM, Law TF, Teixeira P, Wilson ED, Fitzpatrick CR, Jones CD, Dangl JL. 2020. A single bacterial genus maintains root growth in a complex microbiome. Nature 587:103–108.

40. Moore LW, Chilton WS, Canfield ML. 1997. Diversity of opines and opine-catabolizing bacteria isolated from naturally occurring crown gall tumors. Applied and Environmental Microbiology 63:201–207.

41. Beauchamp CJ, Dion P, Kloepper JW, Antoun H. 1991. Physiological Characterization of Opine-Utilizing Rhizobacteria for Traits Related to Plant Growth-Promoting Activity. Plant and Soil 132:273–279.

42. Lin JS, Ma LS, Lai EM. 2013. Systematic Dissection of the Agrobacterium Type VI Secretion System Reveals Machinery and Secreted Components for Subcomplex Formation. PLoS One 8:e67647.

43. Kado CI, and Heskett, M.G.. 1970. Selective media for isolation of *Agrobacterium, Carynebacterium, Erwinia, Pseudomonas*, and *Xanthomonas*. Phytopathology 60:969–976.

44. Massa S, Caruso M, Trovatelli F, Tosques M. 1998. Comparison of plate count agar and R2A medium for enumeration of heterotrophic bacteria in natural mineral water. World Journal of Microbiology & Biotechnology 14:727–730.

45. Ma LS, Lin JS, Lai EM. 2009. An IcmF family protein, ImpLM, is an integral inner membrane protein interacting with ImpKL, and its walker a motif is required for type VI secretion system-mediated Hcp secretion in Agrobacterium tumefaciens. J Bacteriol 191:4316–29.

46. Hawes MC, Smith LY. 1989. Requirement for chemotaxis in pathogenicity of Agrobacterium tumefaciens on roots of soil-grown pea plants. J Bacteriol 171:5668–71.

47. Yakabe LE, Parker SR, Kluepfel DA. 2012. Role of Systemic Agrobacterium tumefaciens Populations in Crown Gall Incidence on the Walnut Hybrid Rootstock ‘Paradox’. Plant Disease 96:1415–1421.

48. McPherson MR, Wang P, Marsh EL, Mitchell RB, Schachtman DP. 2018. Isolation and Analysis of Microbial Communities in Soil, Rhizosphere, and Roots in Perennial Grass Experiments. Jove-Journal of Visualized Experiments doi:ARTN e5793210.3791/57932.

49. Bolyen E, Rideout JR, Dillon MR, Bokulich NA, Abnet CC, Al-Ghalith GA, Alexander H, Alm EJ, Arumugam M, Asnicar F, Bai Y, Bisanz JE, Bittinger K, Brejnrod A, Brislawn CJ, Brown CT, Callahan BJ, Caraballo-Rodriguez AM, Chase J, Cope EK, Da Silva R, Diener C, Dorrestein PC, Douglas GM, Durall DM, Duvallet C, Edwardson CF, Ernst M, Estaki M, Fouquier J, Gauglitz JM, Gibbons SM, Gibson DL, Gonzalez A, Gorlick K, Guo J, Hillmann B, Holmes S, Holste H, Huttenhower C, Huttley GA, Janssen S, Jarmusch AK, Jiang L, Kaehler BD, Kang KB, Keefe CR, Keim P, Kelley ST, Knights D, et al. 2019. Reproducible, interactive, scalable and extensible microbiome data science using QIIME 2. Nat Biotechnol 37:852–857.

50. Callahan BJ, McMurdie PJ, Rosen MJ, Han AW, Johnson AJ, Holmes SP. 2016. DADA2: High-resolution sample inference from Illumina amplicon data. Nat Methods 13:581–3.

51. Rognes T, Flouri T, Nichols B, Quince C, Mahe F. 2016. VSEARCH: a versatile open source tool for metagenomics. PeerJ 4:e2584.

52. Bokulich NA, Kaehler BD, Rideout JR, Dillon M, Bolyen E, Knight R, Huttley GA, Gregory Caporaso J. 2018. Optimizing taxonomic classification of marker-gene amplicon sequences with QIIME 2’s q2-feature-classifier plugin. Microbiome 6:90.

53. Yilmaz P, Parfrey LW, Yarza P, Gerken J, Pruesse E, Quast C, Schweer T, Peplies J, Ludwig W, Glockner FO. 2014. The SILVA and “All-species Living Tree Project (LTP)” taxonomic frameworks. Nucleic Acids Res 42:D643–8.

54. Janssen S, McDonald D, Gonzalez A, Navas-Molina JA, Jiang L, Xu ZZ, Winker K, Kado DM, Orwoll E, Manary M, Mirarab S, Knight R. 2018. Phylogenetic Placement of Exact Amplicon Sequences Improves Associations with Clinical Information. mSystems 3.

55. Matsen FA, Hoffman NG, Gallagher A, Stamatakis A. 2012. A format for phylogenetic placements. PLoS One 7:e31009.

56. Eddy SR. 2011. Accelerated Profile HMM Searches. PLoS Comput Biol 7:e1002195.

57. Matsen FA, Kodner RB, Armbrust EV. 2010. pplacer: linear time maximum-likelihood and Bayesian phylogenetic placement of sequences onto a fixed reference tree. BMC Bioinformatics 11:538.

58. Lozupone C, Knight R. 2005. UniFrac: a new phylogenetic method for comparing microbial communities. Appl Environ Microbiol 71:8228–35.

59. Lin H, Peddada SD. 2020. Analysis of microbial compositions: a review of normalization and differential abundance analysis. NPJ Biofilms Microbiomes 6:60.

60. Mandal S, Van Treuren W, White RA, Eggesbo M, Knight R, Peddada SD. 2015. Analysis of composition of microbiomes: a novel method for studying microbial composition. Microb Ecol Health Dis 26:27663.

61. McMurdie PJ, Holmes S. 2013. phyloseq: an R package for reproducible interactive analysis and graphics of microbiome census data. PLoS One 8:e61217.

62. Anderson MJ. 2001. A new method for non-parametric multivariate analysis of variance. Austral Ecology 26:32–46.

63. Dixon P. 2003. VEGAN, a package of R functions for community ecology. Journal of Vegetation Science 14:927–930.

