## Supplemental figures and tables for "Development of a soil inoculation method coupled with blocker-mediated 16S rRNA gene amplicon sequencing reveals the effect of antibacterial T6SS on agrobacteria tumorigenesis and gallobiome composition"

Figures, S1-S4; Table S1-S8

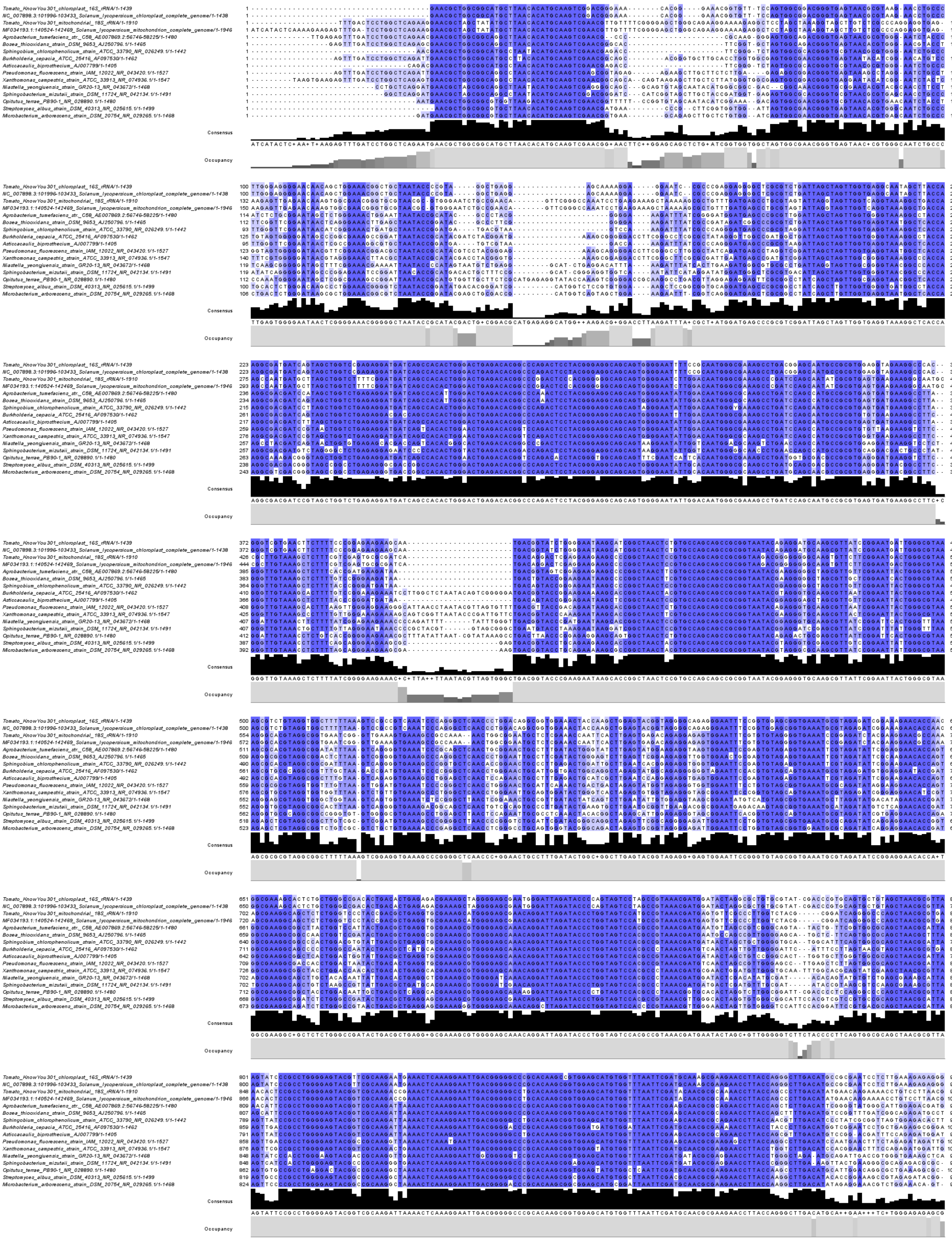
Tomato_KnowYou301_chloroplast_16S_rRNA/1-1439
NC_007898.3:101996-103433_Solanum_lycopersicum_chloroplast_complete_genome/1-1438
Tomato_KnowYou301_mitochondrial_18S_rRNA/1-1910
MF034193.1:140524-142469_Solanum_lycopersicum_mitochondrion_complete_genome/1-1946
Agrobacterium_tumefaciens_str._C58_AE007869.2:56746-58225/1-1480
Bosea_thiooxidans_strain_DSM_9653_AJ250796.1/1-1465
Sphingobium_chlorophenolicum_strain_ATCC_33790_NR_026249.1/1-1442
Burkholderia_cepacia_ATCC_25416_AF097530/1-1462
Asticcacaulis_biprosthecium_AJ007799/1-1405
Pseudomonas_fluorescens_strain_IAM_12022_NR_043420.1/1-1527
Xanthomonas_campestris_strain_ATCC_33913_NR_074936.1/1-1547
Niastella_yeongjuensis_strain_GR20-13_NR_043672/1-1468
Sphingobacterium_mizutaii_strain_DSM_11724_NR_042134.1/1-1491
Opitutus_terrae_PB90-1_NR_028890.1/1-1480
Streptomyces_albus_strain_DSM_40313_NR_025615.1/1-1499
Microbacterium_arborescens_strain_DSM_20754_NR_029265.1/1-1468
Consensus
Occupancy

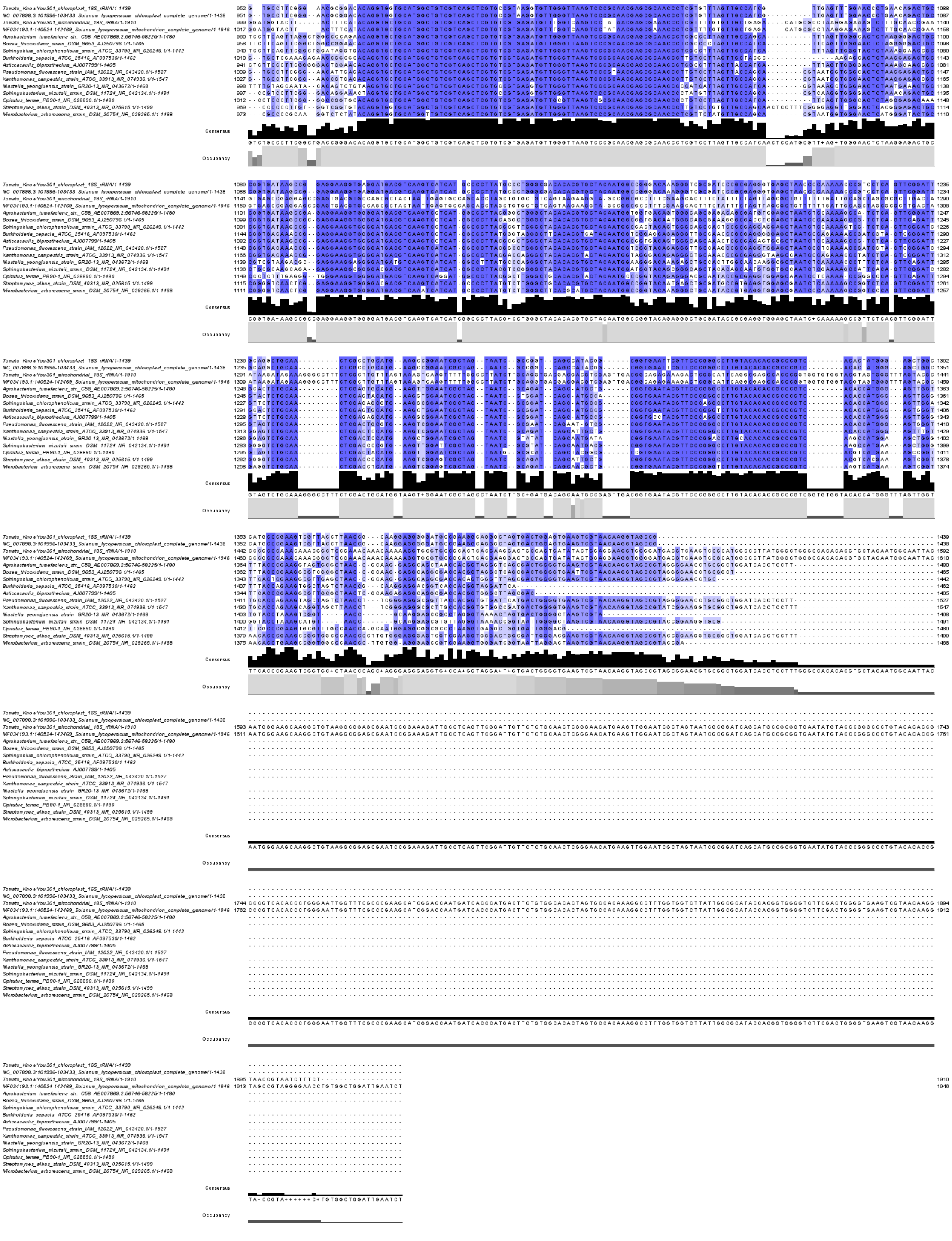

**Figure S1. DNA sequence alignment of 16S rRNA genes of selected bacterial strains and tomato KnowYou 301 chloroplast and mitochondria.**

The 16S rRNA gene sequences of chloroplasts and mitochondria in *Solanum lycopersicum* (tomato) cultivar Known-You 301 were obtained from this study. All of the bacterial 16S rRNA gene sequences were accessed from Reference Sequence (RefSeq) database in National Center of Biotechnology Information (NCBI). The species, strain name, and accession number of 16S rRNA gene were listed on the left of each sequence. The alignment of sequences mentioned above were aligned in MEGAX via ClustalW multiple alignment with gap opening penalty

15.00 and gap extension penalty 6.66 (default). The levels of shaded blue color reflect the degree of identity. The regions of each primer and blocker were underlined. The sequences highlighted by red, black, yellow and green are the annealing region of primer sets for V1-3, V3-4, V5-7 and V6-8 in 16S rRNA gene, respectively. And the gray framed sequences in tomato chloroplast and mitochondrial 16S rRNA genes are the blockers for cognate primers.

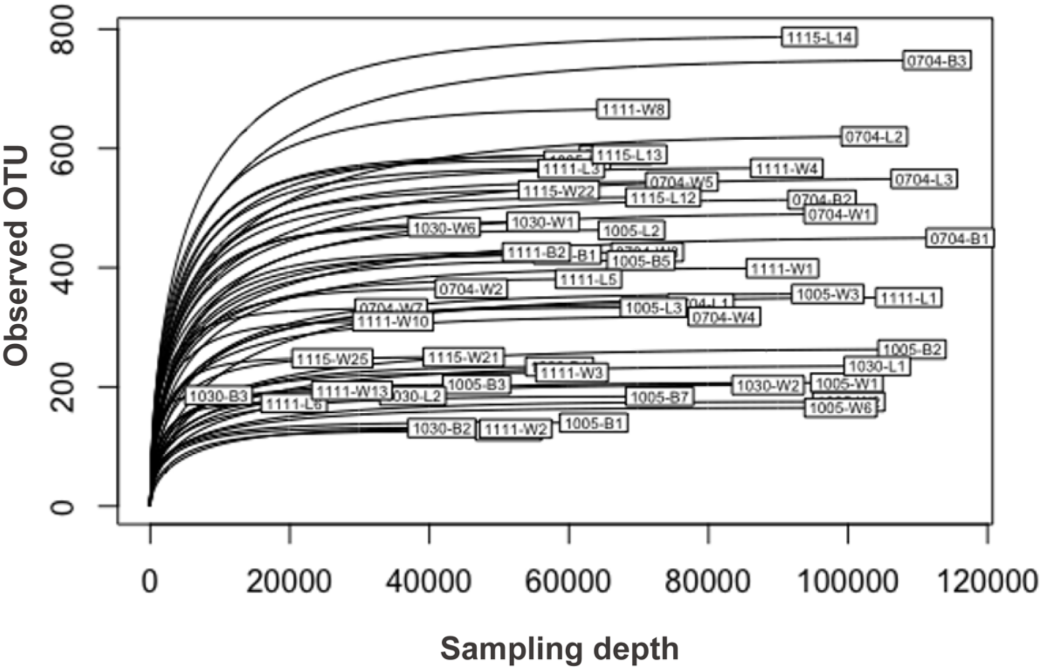


**Figure S2. Alpha rarefaction curves of the observed bacterial OTUs based on amplicon sequencing of 53 gall samples in sequencing run III.**

The rarefaction curves were plotted after clustering DADA2-output ASVs into 99% OTUs. Sample size in X-axis indicates different sub-sampling depth of each dataset; Y-axis indicates the number of observed OTUs under certain sub-sampling depth. Label of each curves indicates the sample ID in metadata.

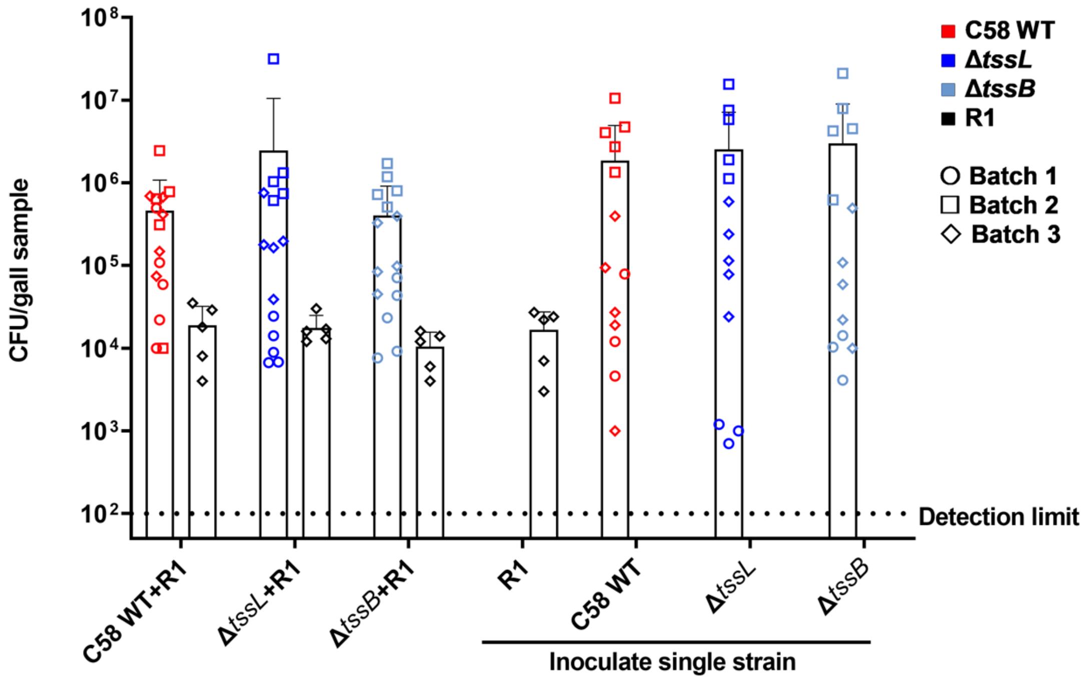


**Figure S3. Competition of *Agrobacterium* and *Sphingomonas* sp. R1 in tomato gall on stem at 28 dpi.** Each of the three *Agrobacterium* strains (i.e., C58 WT, Δ*tssL*, and Δ*tssB*) and R1 were mixed at 1:1 ratio then inoculated on wounded tomato stem. The galls were harvested and homogenized for plating at 28 dpi. *Agrobacterium* strains and *Sphingomonas* sp. R1 were recovered on 523 medium plates containing proper antibiotics. CFU data for agrobacteria are mean ± SD of three independent experiments/batches, each with five seedlings inoculated for each strain. R1 could only be recovered from galls in batch 3. The *p*-value of ANOVA against CFU numbers of C58 for each batch is 0.786, 0.0023 and 0.230; the *p*-value of ANOVA against CFU numbers of R1 was 0.6413. The significance of CFU numbers of C58 in second batch is due to the low CFU counts of Δ*tssL* single inoculation group, which could not be replicated in other two batches.

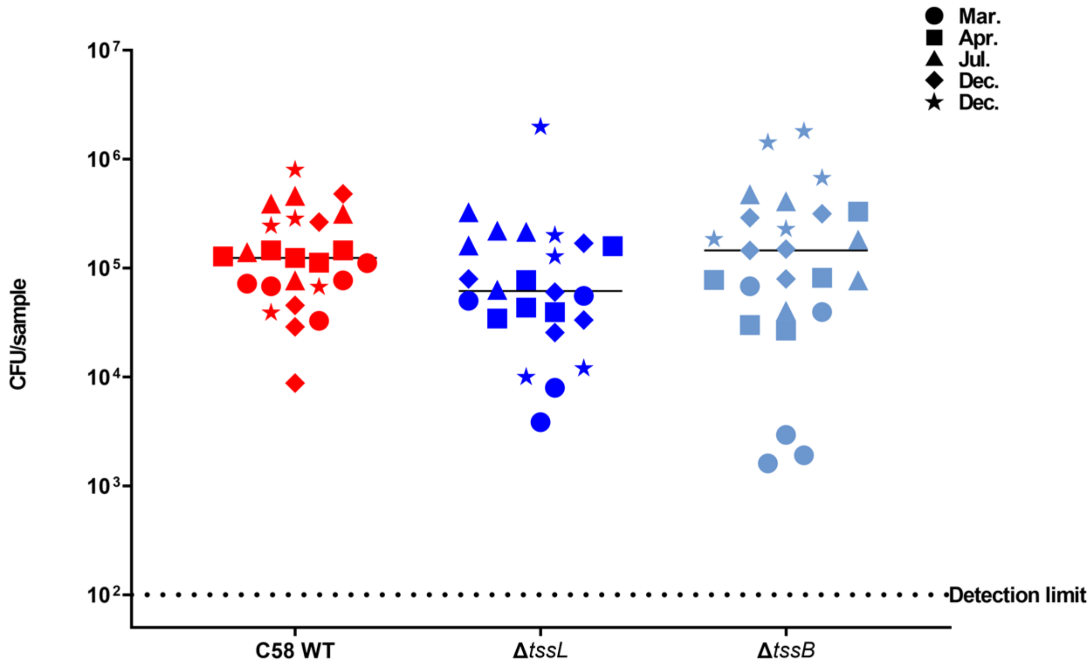


**Figure S4. Agrobacteria colonization on wounded stem segments.**

The colonies of *Agrobacterium* strains C58 WT, Δ*tssL*, and Δ*tssB* recovered from the surface of wounded stem segments following the soil inoculation procedure were plotted. Different symbols indicated the outcome from different batches of colonization assay. *F*-test indicated the variance of Δ*tssL* and Δ*tssB* were significantly different when compared to C58 WT ( $p(F \leq f)$ = 0.00028 and 1.77826E-05, respectively). Brown-Forsythe and Welch ANOVA test indicated no significant difference among means (*p*-value= 0.4847). Line indicates median.

Table S1. 16S rRNA gene primer sets used in this study

| Targeted Region | Name | Sequence (5'-3') | Size of bacterial amplicons (bp) | Size of tomato amplicons (bp)* |
|---|---|---|---|---|
| Round I | | | | |
| V3-V4 | 341F | CCTACGGGNGGCWGCAG | 444 | 396 (cp); 395 (mt) |
| | 785R | GACTACHVGGGTATCTAATCC | | |
| V5-V7 | 799F | AACMGGATTAGATACCCKG | 394 | 376(cp); 742(mt) |
| | 1193R | ACGTCATCCCCACCTTCC | | |
| Round II | | | | |
| V1-V3 | 68F | TNANACATGCAAGTCGRRCG | 425 | 440 (cp); 474(mt) |
| | 518R | WTTACCGCGGCTGCTGG | | |
| V3-V4 | 341F | CCTACGGGNGGCWGCAG | 429 | 408 (cp); 391(mt) |
| | 688R | CGCTTTCGHDCCTCAGYGTCA | 405 | |
| V5-V7 | 819F | GTCCACVCCSTAAACGWTG | 553 | 551 (cp); 915(mt) |
| | 1276R | RCGATTACTAGCGAHTCC | | |
| V6-V8 | 895F | CRCCTGGGGAGTRCRG | 530 | 527 (cp); 894 (mt) |
| | 1391R | GACGGGCGGTGTGTRCA | | |

*cp, chloroplast; mt, mitochondrion

Table S2. Corresponding blockers (3’ modified oligonucleotides with C3 spacer)

| Targeted Region | Name* | Sequence | Tm (℃) |
|---|---|---|---|
| V1 | 68f_blocker_mt | GTCGAACGTTGTTTTCGGGGAG | 62 |
| V1 | 68f_blocker_cp | GTCGGACGGGAAACACG | 61 |
| V4 | 688ra_blocker_mt | CGTCGGTAGGGACCCAGAGAGCT | 69 |
| V4 | 688ra_blocker_cp | TGTCAGTGTCGGCCCAGCAGAGT | 69 |
| V5 | 819f_blocker_mt | ACGATGAGTGTTCGCCCTTG | 61 |
| V5 | 819f_blocker_cp | AAACGATGGATACTAGGTGCTGT | 60 |
| V6 | 895f_blocker_mt | AGTACGGTCGCAAGACCG | 61 |
| V6 | 895f_blocker_cp | GGAGTACGTTCGCAAGAATG | 60 |

* cp, chloroplast; mt, mitochondrion

Table S3. Identification of bacterial isolates from tomato rhizosphere.

| Isolates | Genus | Per. Identity* |
|---|---|---|
| R1 | *Sphingomonas* sp. | 100.00% |
| R3 | *Rhizobium* sp. | 99.43% |
| R4 | *Pseudacidovorax* sp. | 99.81% |
| R5 | *Roseateles* sp. | 100.00% |
| R6 | *Pseudacidovorax* sp. | 99.81% |
| R7 | *Flavobacterium* sp. | 98.86% |
| BR3-1 | *Pseudomonas* sp. | 96.88% |
| BR3-2 | *Acinetobacter* sp. | 99.78% |
| BR3-3 | *Pseudomonas* sp. | 96.98% |
| CKR3-4 | *Pseudomonas* sp. | 97.42% |
| CKR3-5 | *Microbacterium* sp. | 100.00% |
| CKR3-6 | *Pseudomonas* sp. | 99.56% |
| BE3-7 | *Microbacterium* sp. | 100.00% |
| BE3-8 | *Microbacterium* sp. | 100.00% |
| BE3-9 | *Microbacterium* sp. | 100.00% |
| BE5-10 | *Microbacterium* sp. | 100.00% |

* The partial 16S rRNA genes were amplified by V5-V7 primer set and the sequences were blast against 16S ribosomal RNA sequences database in NCBI.

**Table S4.** Bacterial strains and plasmids used in this study

| Strain/plasmid | Characteristics | EML No. | Reference/ Source |
|---|---|---|---|
| *Escherichia coli* DH10B | Host for DNA cloning | 455 | Invitrogen |
| *Agrobacterium* C58 | Wild-type, virulent strain containing nonpaline-type Ti-plasimid, pTiC58 | 530 | Eugene Nester [1] |
| | Δ*tssL*, deletion mutant of *tssL* gene (Atu4333) | 1073 | [2] |
| | Δ*tssB*, deletion mutant of *tssB* gene (Atu4342) | 1109 | [2] |
| | $Gm^R$-GFP knocked-in chromosome | 4094 | This study |
| | Δ*tssL*, $Gm^R$-GFP knocked-in chromosome | 4097 | This study |
| | Δ*tssB*, $Gm^R$-GFP knocked-in chromosome | 4100 | This study |
| *Sphingomonas sp.* R1 | Wild type, isolated from tomato rhizosphere | 4107 | This study |
| pJQ-COM | $Gm^R$, for generating DNA knock-in adjacent to gene *ActC* in C58 | 2236 | [3] |
| pJQ-com-$Gm^r$ GFP | $Gm^R$, for generate $Gm^R$ and GFP knocked-in C58 | 4091 | This study |
| pRL662::GFP (S65T) | $Gm^R$, constitutively expressing GFP (S56T) | 3375 | This study |
| pBBR1MCS2-mCherry | $Km^R$, constitutively expressing mCherry | 3022 | Stanton B. Gelvin |

Table S5. Primers used for plasmid construction

| Name | Sequence | Reference |
|---|---|---|
| BclI-GFP-GmR-F-25 | AAAAATGATCAGTGAGCGCGCGTAATA CGACTCAC | This study |
| BclI-GFP-GmR-R-25 | AAAAATGATCAGGGTACCGAGCTCGAA TTGACATAAG | This study |
| 3'sacB-5'GmR pJQ200 F | TGCGCCAAGCTTCCTGCTGAACATC | This study |
| 3'sacB-5'GmR pJQ200 R | TTGAGCAGCCGCGTAGTGAGATCTA | This study |

Table S6. Condition of 16S rRNA gene amplification

| 1st PCR Amplification v1-3 | | | | | | | | | | 2nd Library Amplification | | | | |
|---|---|---|---|---|---|---|---|---|---|---|---|---|---|---|
| **Recipe A** | | **Program** | | | **Recipe B** | | **Program** | | | **Recipe** | | **Program** | | |
| input 25 ng | 6.5 | 95℃ | 5 min | 1 cycle | input 25 ng | 6.5 | 95℃ | 5 min | 1 cycle | 1st purified PCR products | 20 | 98℃ | 2 min | 1 cycle |
| H2O | | 98℃ | 30 sec | | H2O | | 98℃ | 30 sec | | H2O | | 98℃ | **30** sec | |
| 2X **Kapa HiFi PCR mix** | 12.5 | **55℃** | 30 sec | 12cycle | 2X **Kapa HiFi PCR mix** | 12.5 | **55℃** | 30 sec | 15cycle | 2X **Kapa HiFi PCR mix** | 25 | 60℃ | 30 sec | **5** cycle |
| 68f primer 10uM (final 0.2uM) | 0.5 | 72℃ | 60 sec | | 68f primer 10uM (final 0.2uM) | 0.5 | 72℃ | 60 sec | | Truseq-HT-D5xx_F (10uM) | 2.5 | 72℃ | 60 sec | |
| 518r primer 10uM (final 0.2uM) | 0.5 | 72℃ | 5 min | 1 cycle | 518r primer 10uM (final 0.2uM | 0.5 | 72℃ | 5 min | 1 cycle | Truseq-HT-D7xx_R (10uM) | 2.5 | 72℃ | 5 min | 1 cycle |
| H2O | 5 | 4℃ | ∞ | 1 cycle | 68f_blocking_mito 20uM | 2.5 | 4℃ | ∞ | 1 cycle | Total | 50 | 10℃ | ∞ | 1 cycle |
| | | | | | 518r_blocking_chloro_20uM | 2.5 | | | | | | | | |
| Total | 25 | | | | Total | 25 | | | | 25uL/tube | | | | |
| Beads purificatin, 1:1 | | | | | Beads purificatin, 1:1 | | | | | Beads purificatin, 1:1 | | | | |

| 1st PCR Amplification v3-4 | | | | | | | | | | 2nd Library Amplification | | | | |
|---|---|---|---|---|---|---|---|---|---|---|---|---|---|---|
| **Recipe A** | | **Program** | | | **Recipe B** | | **Program** | | | **Recipe** | | **Program** | | |
| input 25 ng | 6.5 | 95℃ | 5 min | 1 cycle | input 25 ng | 6.5 | 95℃ | 5 min | 1 cycle | 1st purified PCR products | 20 | 98℃ | 2 min | 1 cycle |
| H2O | | 98℃ | 30 sec | | H2O | | 98℃ | 30 sec | | H2O | | 98℃ | **30** sec | 5 cycle (A)<br>10 cycle (B) |
| 2X **Kapa HiFi PCR mix** | 12.5 | **56℃** | 30 sec | 18cycle | 2X **Kapa HiFi PCR mix** | 12.5 | **56℃** | 30 sec | 18 cycle | 2X **Kapa HiFi PCR mix** | 25 | 60℃ | 30 sec | |
| 341f primer 10uM (final 0.2uM) | 0.5 | 72℃ | 60 sec | | 341f primer 10uM (final 0.2uM | 0.5 | 72℃ | 60 sec | | Truseq-HT-D5xx_F (10uM) | 2.5 | 72℃ | 60 sec | |
| 688ar primer 10uM (final 0.2uM | 0.5 | 72℃ | 5 min | 1 cycle | 688ar primer 10uM (final 0.2ul | 0.5 | 72℃ | 5 min | 1 cycle | Truseq-HT-D7xx_R (10uM) | 2.5 | 72℃ | 5 min | 1 cycle |
| H2O | 5 | 4℃ | ∞ | 1 cycle | 688ar_blocking_mito 20uM | 2.5 | 4℃ | ∞ | 1 cycle | Total | 50 | 10℃ | ∞ | 1 cycle |
| | | | | | 688ar_blocking_chloro_20uM | 2.5 | | | | | | | | |
| Total | 25 | | | | Total | 25 | | | | 25uL/tube | | | | |
| Beads purificatin, 1:1 | | | | | Beads purificatin, 1:1 | | | | | Beads purificatin, 1:1 | | | | |

| 1st PCR Amplification v5-7 | | | | | | | | | | 2nd Library Amplification | | | | |
|---|---|---|---|---|---|---|---|---|---|---|---|---|---|---|
| **Recipe A** | | **Program** | | | **Recipe B** | | **Program** | | | **Recipe** | | **Program** | | |
| input 25 ng | 6.5 | 95℃ | 5 min | 1 cycle | input 25 ng | 6.5 | 95℃ | 5 min | 1 cycle | 1st purified PCR products | 20 | 98℃ | 2 min | 1 cycle |
| H2O | | 98℃ | 30 sec | | H2O | | 98℃ | 30 sec | | H2O | | 98℃ | **30** sec | |
| 2X **Kapa HiFi PCR mix** | 12.5 | **56℃** | 30 sec | 20cycle | 2X **Kapa HiFi PCR mix** | 12.5 | **56℃** | 30 sec | 24cycle | 2X **Kapa HiFi PCR mix** | 25 | 60℃ | 30 sec | **5** cycle |
| 819f primer 10uM (final 0.2uM) | 0.5 | 72℃ | 60 sec | | 819f primer 10uM (final 0.2uM | 0.5 | 72℃ | 60 sec | | Truseq-HT-D5xx_F (10uM) | 2.5 | 72℃ | 60 sec | |
| 1276r primer 10uM (final 0.2uM | 0.5 | 72℃ | 5 min | 1 cycle | 1276r primer 10uM (final 0.2ul | 0.5 | 72℃ | 5 min | 1 cycle | Truseq-HT-D7xx_R (10uM) | 2.5 | 72℃ | 5 min | 1 cycle |
| H2O | 5 | 4℃ | ∞ | 1 cycle | 819f_blocking_mito 20uM | 2.5 | 4℃ | ∞ | 1 cycle | Total | 50 | 10℃ | forever | 1 cycle |
| | | | | | 819f_blocking_chloro_20uM | 2.5 | | | | | | | | |
| Total | 25 | | | | Total | 25 | | | | 25uL/tube | | | | |
| Beads purificatin, 1:1 | | | | | Beads purificatin, 1:1 | | | | | Beads purificatin, 1:1 | | | | |

| 1st PCR Amplification v6-8 | | | | | | | | | | 2nd Library Amplification | | | | |
|---|---|---|---|---|---|---|---|---|---|---|---|---|---|---|
| **Recipe A** | | **Program** | | | **Recipe B** | | **Program** | | | **Recipe** | | **Program** | | |
| input 25 ng | 6.5 | 95℃ | 5 min | 1 cycle | input 25 ng | 6.5 | 95℃ | 5 min | 1 cycle | 1st purified PCR products | 20 | 98℃ | 2 min | 1 cycle |
| H2O | | 98℃ | 30 sec | | H2O | | 98℃ | 30 sec | | H2O | | 98℃ | **30** sec | |
| 2X **Kapa HiFi PCR mix** | 12.5 | **50℃** | 30 sec | 11cycle | 2X **Kapa HiFi PCR mix** | 12.5 | **50℃** | 30 sec | 12cycle | 2X **Kapa HiFi PCR mix** | 25 | 60℃ | 30 sec | **5** cycle |
| 895f primer 10uM (final 0.2uM) | 0.5 | 72℃ | 60 sec | | 895f primer 10uM (final 0.2uM | 0.5 | 72℃ | 60 sec | | Truseq-HT-D5xx_F (10uM) | 2.5 | 72℃ | 60 sec | |
| 1391r primer 10uM (final 0.2uM | 0.5 | 72℃ | 5 min | 1 cycle | 1391r primer 10uM (final 0.2ul | 0.5 | 72℃ | 5 min | 1 cycle | Truseq-HT-D7xx_R (10uM) | 2.5 | 72℃ | 5 min | 1 cycle |
| H2O | 5 | 4℃ | ∞ | 1 cycle | 895f_blocking_mito 20uM | 2.5 | 4℃ | ∞ | 1 cycle | Total | 50 | 10℃ | forever | 1 cycle |
| | | | | | 895f_blocking_chloro_20uM | 2.5 | | | | | | | | |
| Total | 25 | | | | Total | 25 | | | | 25uL/tube | | | | |
| Beads purificatin, 1:1 | | | | | Beads purificatin, 1:1 | | | | | Beads purificatin, 1:1 | | | | |

Table S7. Metadata of harvested crown galls

| | | | | | Harvest time | | | |
|---|---|---|---|---|---|---|---|---|
| #Sample ID | Strain* | Year of inoculation | Month of inoculation | Day of inoculation | Year | Month | Day | Weight (g) |
| 1115-11L | Δ*tssL* | 2018 | 11 | 15 | 2019 | 1 | 18 | 1.80 |
| 1115-12L[a] | Δ*tssL* | 2018 | 11 | 15 | 2019 | 1 | 18 | 0.81 |
| 1115-13L[a] | Δ*tssL* | 2018 | 11 | 15 | 2019 | 1 | 18 | 0.62 |
| 1115-14L[a] | Δ*tssL* | 2018 | 11 | 15 | 2019 | 1 | 18 | 0.96 |
| 1115-15L | Δ*tssL* | 2018 | 11 | 15 | 2019 | 1 | 18 | 0.40 |
| 1115-16L | Δ*tssL* | 2018 | 11 | 15 | 2019 | 1 | 18 | 0.40 |
| 1115-17L | Δ*tssL* | 2018 | 11 | 15 | 2019 | 1 | 18 | 0.07 |
| 1115-18L | Δ*tssL* | 2018 | 11 | 15 | 2019 | 1 | 18 | 0.16 |
| 1115-19W | WT | 2018 | 11 | 15 | 2019 | 1 | 18 | 2.00 |
| 1115-20W | WT | 2018 | 11 | 15 | 2019 | 1 | 18 | 1.27 |
| 1115-21W[b] | WT | 2018 | 11 | 15 | 2019 | 1 | 18 | 0.56 |
| 1115-22W[b] | WT | 2018 | 11 | 15 | 2019 | 1 | 18 | 0.56 |
| 1115-23W | WT | 2018 | 11 | 15 | 2019 | 1 | 18 | 0.42 |
| 1115-24W | WT | 2018 | 11 | 15 | 2019 | 1 | 18 | 0.37 |
| 1115-25W[b] | WT | 2018 | 11 | 15 | 2019 | 1 | 18 | 0.82 |
| 1115-26W | WT | 2018 | 11 | 15 | 2019 | 1 | 18 | 0.22 |
| 1115-27W | WT | 2018 | 11 | 15 | 2019 | 1 | 18 | 0.10 |
| 1115-28W | WT | 2018 | 11 | 15 | 2019 | 1 | 18 | 0.08 |
| 1115-29W | WT | 2018 | 11 | 15 | 2019 | 1 | 18 | 0.05 |
| 1115-30W | WT | 2018 | 11 | 15 | 2019 | 1 | 18 | 0.02 |
| 0704-1W | WT | 2019 | 7 | 4 | 2019 | 9 | 2 | 0.12 |
| 0704-2W | WT | 2019 | 7 | 4 | 2019 | 9 | 2 | 0.12 |
| 0704-3W | WT | 2019 | 7 | 4 | 2019 | 9 | 2 | 0.09 |
| 0704-4W | WT | 2019 | 7 | 4 | 2019 | 9 | 2 | 0.06 |
| 0704-5W | WT | 2019 | 7 | 4 | 2019 | 9 | 2 | 0.11 |
| 0704-6W | WT | 2019 | 7 | 4 | 2019 | 9 | 2 | 0.14 |
| 0704-7W | WT | 2019 | 7 | 4 | 2019 | 9 | 2 | 0.27 |
| 0704-1L | Δ*tssL* | 2019 | 7 | 4 | 2019 | 9 | 2 | 0.10 |
| 0704-2L | Δ*tssL* | 2019 | 7 | 4 | 2019 | 9 | 2 | 0.34 |
| 0704-3L | Δ*tssL* | 2019 | 7 | 4 | 2019 | 9 | 2 | 0.12 |
| 0704-4L | Δ*tssL* | 2019 | 7 | 4 | 2019 | 9 | 2 | 0.07 |
| 0704-5L | Δ*tssL* | 2019 | 7 | 4 | 2019 | 9 | 2 | 0.06 |
| 0704-6L | Δ*tssL* | 2019 | 7 | 4 | 2019 | 9 | 2 | 0.10 |
| 0704-7L | Δ*tssL* | 2019 | 7 | 4 | 2019 | 9 | 2 | 0.13 |

| | | | | | Harvest time | | | |
|---|---|---|---|---|---|---|---|---|
| #Sample ID | Strain* | Year of inoculation | Month of inoculation | Day of inoculation | Year | Month | Day | Weight (g) |
| 0704-1B | Δ*tssB* | 2019 | 7 | 4 | 2019 | 9 | 2 | 0.11 |
| 0704-2B | Δ*tssB* | 2019 | 7 | 4 | 2019 | 9 | 2 | 0.28 |
| 0704-3B | Δ*tssB* | 2019 | 7 | 4 | 2019 | 9 | 2 | 0.07 |
| 0704-4B | Δ*tssB* | 2019 | 7 | 4 | 2019 | 9 | 2 | 0.22 |
| 0704-5B | Δ*tssB* | 2019 | 7 | 4 | 2019 | 9 | 2 | 0.06 |
| 0722-1W | WT | 2019 | 7 | 22 | 2019 | 9 | 28 | 0.03 |
| 0722-2W | WT | 2019 | 7 | 22 | 2019 | 9 | 28 | 0.04 |
| 0722-3W | WT | 2019 | 7 | 22 | 2019 | 9 | 28 | 0.07 |
| 0722-4W | WT | 2019 | 7 | 22 | 2019 | 9 | 28 | 0.05 |
| 0722-5W | WT | 2019 | 7 | 22 | 2019 | 9 | 28 | 0.13 |
| 0722-6W | WT | 2019 | 7 | 22 | 2019 | 9 | 28 | 0.05 |
| 0722-1B | Δ*tssB* | 2019 | 7 | 22 | 2019 | 9 | 28 | 0.01 |
| 0722-2B | Δ*tssB* | 2019 | 7 | 22 | 2019 | 9 | 28 | 0.01 |
| 0729-1W | WT | 2019 | 7 | 29 | 2019 | 9 | 29 | 0.03 |
| 0729-2W | WT | 2019 | 7 | 29 | 2019 | 9 | 29 | 0.05 |
| 0729-3W | WT | 2019 | 7 | 29 | 2019 | 9 | 29 | 0.04 |
| 0729-4W | WT | 2019 | 7 | 29 | 2019 | 9 | 29 | 0.05 |
| 0729-5W | WT | 2019 | 7 | 29 | 2019 | 9 | 29 | 0.04 |
| 0729-6W | WT | 2019 | 7 | 29 | 2019 | 9 | 29 | 0.01 |
| 0729-7W | WT | 2019 | 7 | 29 | 2019 | 9 | 29 | 0.05 |
| 0729-8W | WT | 2019 | 7 | 29 | 2019 | 9 | 29 | 0.08 |
| 0729-9W | WT | 2019 | 7 | 29 | 2019 | 9 | 29 | 0.03 |
| 0729-1L | Δ*tssL* | 2019 | 7 | 29 | 2019 | 9 | 29 | 0.01 |
| 0729-2L | Δ*tssL* | 2019 | 7 | 29 | 2019 | 9 | 29 | 0.05 |
| 0729-3L | Δ*tssL* | 2019 | 7 | 29 | 2019 | 9 | 29 | 0.08 |
| 0729-4L | Δ*tssL* | 2019 | 7 | 29 | 2019 | 9 | 29 | 0.03 |
| 0729-1B | ΔtssB | 2019 | 7 | 29 | 2019 | 9 | 29 | 0.09 |
| 0729-2B | ΔtssB | 2019 | 7 | 29 | 2019 | 9 | 29 | 0.07 |
| 1005-1W | WT | 2019 | 10 | 5 | 2019 | 12 | 13 | 0.27 |
| 1005-2W | WT | 2019 | 10 | 5 | 2019 | 12 | 13 | 0.16 |
| 1005-3W | WT | 2019 | 10 | 5 | 2019 | 12 | 13 | 0.09 |
| 1005-4W | WT | 2019 | 10 | 5 | 2019 | 12 | 13 | 0.04 |
| 1005-5W | WT | 2019 | 10 | 5 | 2019 | 12 | 13 | 0.01 |
| 1005-6W | WT | 2019 | 10 | 5 | 2019 | 12 | 13 | 0.09 |
| 1005-7W | WT | 2019 | 10 | 5 | 2019 | 12 | 13 | 0.01 |

| | | | | | Harvest time | | | |
|---|---|---|---|---|---|---|---|---|
| #Sample ID | Strain* | Year of inoculation | Month of inoculation | Day of inoculation | Year | Month | Day | Weight (g) |
| 1005-1L | Δ*tssL* | 2019 | 10 | 5 | 2019 | 12 | 13 | 0.23 |
| 1005-2L | Δ*tssL* | 2019 | 10 | 5 | 2019 | 12 | 13 | 0.32 |
| 1005-3L | Δ*tssL* | 2019 | 10 | 5 | 2019 | 12 | 13 | 0.09 |
| 1005-4L | Δ*tssL* | 2019 | 10 | 5 | 2019 | 12 | 13 | 0.04 |
| 1005-5L | Δ*tssL* | 2019 | 10 | 5 | 2019 | 12 | 13 | 0.01 |
| 1005-1B | Δ*tssB* | 2019 | 10 | 5 | 2019 | 12 | 13 | 0.23 |
| 1005-2B | Δ*tssB* | 2019 | 10 | 5 | 2019 | 12 | 13 | 0.56 |
| 1005-3B | Δ*tssB* | 2019 | 10 | 5 | 2019 | 12 | 13 | 0.16 |
| 1005-4B | Δ*tssB* | 2019 | 10 | 5 | 2019 | 12 | 13 | 0.05 |
| 1005-5B | Δ*tssB* | 2019 | 10 | 5 | 2019 | 12 | 13 | 0.06 |
| 1005-6B | Δ*tssB* | 2019 | 10 | 5 | 2019 | 12 | 13 | 0.03 |
| 1005-7B | Δ*tssB* | 2019 | 10 | 5 | 2019 | 12 | 13 | 0.06 |
| 1030-1W | WT | 2019 | 10 | 30 | 2020 | 1 | 9 | 0.34 |
| 1030-2W | WT | 2019 | 10 | 30 | 2020 | 1 | 9 | 0.12 |
| 1030-3W | WT | 2019 | 10 | 30 | 2020 | 1 | 9 | 0.05 |
| 1030-4W | WT | 2019 | 10 | 30 | 2020 | 1 | 9 | 0.03 |
| 1030-5W | WT | 2019 | 10 | 30 | 2020 | 1 | 9 | 0.05 |
| 1030-6W | WT | 2019 | 10 | 30 | 2020 | 1 | 9 | 0.11 |
| 1030-7W | WT | 2019 | 10 | 30 | 2020 | 1 | 9 | 0.02 |
| 1030-8W | WT | 2019 | 10 | 30 | 2020 | 1 | 9 | 0.01 |
| 1030-1L | Δ*tssL* | 2019 | 10 | 30 | 2020 | 1 | 9 | 0.50 |
| 1030-2L | Δ*tssL* | 2019 | 10 | 30 | 2020 | 1 | 9 | 0.07 |
| 1030-3L | Δ*tssL* | 2019 | 10 | 30 | 2020 | 1 | 9 | 0.03 |
| 1030-4L | Δ*tssL* | 2019 | 10 | 30 | 2020 | 1 | 9 | 0.01 |
| 1030-1B | Δ*tssB* | 2019 | 10 | 30 | 2020 | 1 | 9 | 0.17 |
| 1030-2B | Δ*tssB* | 2019 | 10 | 30 | 2020 | 1 | 9 | 0.06 |
| 1030-3B | Δ*tssB* | 2019 | 10 | 30 | 2020 | 1 | 9 | 0.08 |
| 1030-4B | Δ*tssB* | 2019 | 10 | 30 | 2020 | 1 | 9 | 0.00 |
| 1030-5B | Δ*tssB* | 2019 | 10 | 30 | 2020 | 1 | 9 | 0.02 |
| 1030-6B | Δ*tssB* | 2019 | 10 | 30 | 2020 | 1 | 9 | 0.01 |
| 1111-1W | WT | 2019 | 11 | 11 | 2020 | 1 | 14 | 0.26 |
| 1111-2W | WT | 2019 | 11 | 11 | 2020 | 1 | 14 | 0.19 |
| 1111-2.1W | WT | 2019 | 11 | 11 | 2020 | 1 | 14 | 0.04 |
| 1111-3W | WT | 2019 | 11 | 11 | 2020 | 1 | 14 | 0.25 |
| 1111-3.1W | WT | 2019 | 11 | 11 | 2020 | 1 | 14 | 0.04 |

| | | | | | Harvest time | | | |
|---|---|---|---|---|---|---|---|---|
| #Sample ID | Strain* | Year of inoculation | Month of inoculation | Day of inoculation | Year | Month | Day | Weight (g) |
| 1111-4W | WT | 2019 | 11 | 11 | 2020 | 1 | 14 | 0.13 |
| 1111-5W | WT | 2019 | 11 | 11 | 2020 | 1 | 14 | 0.01 |
| 1111-6W | WT | 2019 | 11 | 11 | 2020 | 1 | 14 | 0.04 |
| 1111-7W | WT | 2019 | 11 | 11 | 2020 | 1 | 14 | 0.05 |
| 1111-8W | WT | 2019 | 11 | 11 | 2020 | 1 | 14 | 0.11 |
| 1111-8.1W | WT | 2019 | 11 | 11 | 2020 | 1 | 14 | 0.09 |
| 1111-8.2W | WT | 2019 | 11 | 11 | 2020 | 1 | 14 | 0.09 |
| 1111-9W | WT | 2019 | 11 | 11 | 2020 | 1 | 14 | 0.02 |
| 1111-10W | WT | 2019 | 11 | 11 | 2020 | 1 | 14 | 0.07 |
| 1111-11W | WT | 2019 | 11 | 11 | 2020 | 1 | 14 | 0.02 |
| 1111-12W | WT | 2019 | 11 | 11 | 2020 | 1 | 14 | 0.04 |
| 1111-13W | WT | 2019 | 11 | 11 | 2020 | 1 | 14 | 0.27 |
| 1111-13.1W | WT | 2019 | 11 | 11 | 2020 | 1 | 14 | 0.18 |
| 1111-13.2W | WT | 2019 | 11 | 11 | 2020 | 1 | 14 | 0.12 |
| 1111-14W | WT | 2019 | 11 | 11 | 2020 | 1 | 14 | 0.04 |
| 1111-15W | WT | 2019 | 11 | 11 | 2020 | 1 | 14 | 0.01 |
| 1111-1L | Δ*tssL* | 2019 | 11 | 11 | 2020 | 1 | 14 | 0.21 |
| 1111-2L | Δ*tssL* | 2019 | 11 | 11 | 2020 | 1 | 14 | 0.04 |
| 1111-2.1L | Δ*tssL* | 2019 | 11 | 11 | 2020 | 1 | 14 | 0.02 |
| 1111-3L | Δ*tssL* | 2019 | 11 | 11 | 2020 | 1 | 14 | 0.07 |
| 1111-3.1L | Δ*tssL* | 2019 | 11 | 11 | 2020 | 1 | 14 | 0.03 |
| 1111-4L | Δ*tssL* | 2019 | 11 | 11 | 2020 | 1 | 14 | 0.03 |
| 1111-5L | Δ*tssL* | 2019 | 11 | 11 | 2020 | 1 | 14 | 0.08 |
| 1111-6L | Δ*tssL* | 2019 | 11 | 11 | 2020 | 1 | 14 | 0.08 |
| 1111-7L | Δ*tssL* | 2019 | 11 | 11 | 2020 | 1 | 14 | 0.14 |
| 1111-8L | Δ*tssL* | 2019 | 11 | 11 | 2020 | 1 | 14 | 0.01 |
| 1111-9L | Δ*tssL* | 2019 | 11 | 11 | 2020 | 1 | 14 | 0.03 |
| 1111-1B | Δ*tssB* | 2019 | 11 | 11 | 2020 | 1 | 14 | 0.18 |
| 1111-2B | Δ*tssB* | 2019 | 11 | 11 | 2020 | 1 | 14 | 0.09 |
| 1111-3B | Δ*tssB* | 2019 | 11 | 11 | 2020 | 1 | 14 | 0.03 |
| 1111-4B | Δ*tssB* | 2019 | 11 | 11 | 2020 | 1 | 14 | 0.03 |
| 1111-5B | Δ*tssB* | 2019 | 11 | 11 | 2020 | 1 | 14 | 0.02 |
| 1111-6B | Δ*tssB* | 2019 | 11 | 11 | 2020 | 1 | 14 | 0.03 |
| 1111-7B | Δ*tssB* | 2019 | 11 | 11 | 2020 | 1 | 14 | 0.01 |
| 1111-8B | Δ*tssB* | 2019 | 11 | 11 | 2020 | 1 | 14 | 0.01 |

* *Agrobacterium* C58, [a] Used in Round I sequencing, [b] Used in both Round I and II sequencing.

Table S8. Crown gall metadata used for analysis of amplicon sequencing round III

| #Sample ID | Strain | Year_of_ inoculation | Month_of_ inoculation | Weight | Season | Trial | Batch |
|---|---|---|---|---|---|---|---|
| #q2:types | category | category | category | numeric | category | category | category |
| 1115-L12 | Δ*tssL* | 2018 | Nov | 0.81 | Winter | 181115 | Batch 1 |
| 1115-L13 | Δ*tssL* | 2018 | Nov | 0.62 | Winter | 181115 | Batch 1 |
| 1115-L14 | Δ*tssL* | 2018 | Nov | 0.96 | Winter | 181115 | Batch 1 |
| 1115-W21 | WT | 2018 | Nov | 0.56 | Winter | 181115 | Batch 1 |
| 1115-W22 | WT | 2018 | Nov | 0.56 | Winter | 181115 | Batch 1 |
| 1115-W25 | WT | 2018 | Nov | 0.82 | Winter | 181115 | Batch 1 |
| 0704-W1 | WT | 2019 | Jul | 0.12 | Summer | 190704 | Batch 4 |
| 0704-W2 | WT | 2019 | Jul | 0.12 | Summer | 190704 | Batch 4 |
| 0704-W3 | WT | 2019 | Jul | 0.09 | Summer | 190704 | Batch 4 |
| 0704-W4 | WT | 2019 | Jul | 0.06 | Summer | 190704 | Batch 4 |
| 0704-W5 | WT | 2019 | Jul | 0.11 | Summer | 190704 | Batch 4 |
| 0704-W6 | WT | 2019 | Jul | 0.14 | Summer | 190704 | Batch 4 |
| 0704-W7 | WT | 2019 | Jul | 0.27 | Summer | 190704 | Batch 4 |
| 0704-L1 | Δ*tssL* | 2019 | Jul | 0.1 | Summer | 190704 | Batch 4 |
| 0704-L2 | Δ*tssL* | 2019 | Jul | 0.34 | Summer | 190704 | Batch 4 |
| 0704-L3 | Δ*tssL* | 2019 | Jul | 0.12 | Summer | 190704 | Batch 4 |
| 0704-B1 | Δ*tssB* | 2019 | Jul | 0.11 | Summer | 190704 | Batch 4 |
| 0704-B2 | Δ*tssB* | 2019 | Jul | 0.28 | Summer | 190704 | Batch 4 |
| 0704-B3 | Δ*tssB* | 2019 | Jul | 0.07 | Summer | 190704 | Batch 4 |
| 1005-W1 | WT | 2019 | Oct | 0.27 | Summer | 191005 | Batch 7 |
| 1005-W2 | WT | 2019 | Oct | 0.16 | Summer | 191005 | Batch 7 |
| 1005-W3 | WT | 2019 | Oct | 0.09 | Summer | 191005 | Batch 7 |
| 1005-W6 | WT | 2019 | Oct | 0.09 | Summer | 191005 | Batch 7 |
| 1005-L1 | Δ*tssL* | 2019 | Oct | 0.23 | Summer | 191005 | Batch 7 |
| 1005-L2 | Δ*tssL* | 2019 | Oct | 0.32 | Summer | 191005 | Batch 7 |
| 1005-L3 | Δ*tssL* | 2019 | Oct | 0.09 | Summer | 191005 | Batch 7 |
| 1005-B1 | Δ*tssB* | 2019 | Oct | 0.23 | Summer | 191005 | Batch 7 |
| 1005-B2 | Δ*tssB* | 2019 | Oct | 0.56 | Summer | 191005 | Batch 7 |
| 1005-B3 | Δ*tssB* | 2019 | Oct | 0.16 | Summer | 191005 | Batch 7 |
| 1005-B5 | Δ*tssB* | 2019 | Oct | 0.06 | Summer | 191005 | Batch 7 |
| 1005-B7 | Δ*tssB* | 2019 | Oct | 0.06 | Summer | 191005 | Batch 7 |
| 1030-W1 | WT | 2019 | Oct | 0.34 | Winter | 191030 | Batch 8 |
| 1030-W2 | WT | 2019 | Oct | 0.12 | Winter | 191030 | Batch 8 |
| 1030-W6 | WT | 2019 | Oct | 0.11 | Winter | 191030 | Batch 8 |

| #Sample ID | Strain | Year_of_ inoculation | Month_of_ inoculation | Weight | Season | Trial | Batch |
|---|---|---|---|---|---|---|---|
| 1030-L1 | Δ*tssL* | 2019 | Oct | 0.5 | Winter | 191030 | Batch 8 |
| 1030-L2 | Δ*tssL* | 2019 | Oct | 0.07 | Winter | 191030 | Batch 8 |
| 1030-B1 | Δ*tssL* | 2019 | Oct | 0.17 | Winter | 191030 | Batch 8 |
| 1030-B2 | Δ*tssL* | 2019 | Oct | 0.06 | Winter | 191030 | Batch 8 |
| 1030-B3 | Δ*tssL* | 2019 | Oct | 0.08 | Winter | 191030 | Batch 8 |
| 1111-W1 | WT | 2019 | Nov | 0.26 | Winter | 191111 | Batch 9 |
| 1111-W2 | WT | 2019 | Nov | 0.19 | Winter | 191111 | Batch 9 |
| 1111-W3 | WT | 2019 | Nov | 0.25 | Winter | 191111 | Batch 9 |
| 1111-W4 | WT | 2019 | Nov | 0.13 | Winter | 191111 | Batch 9 |
| 1111-W8 | WT | 2019 | Nov | 0.11 | Winter | 191111 | Batch 9 |
| 1111-W10 | WT | 2019 | Nov | 0.07 | Winter | 191111 | Batch 9 |
| 1111-W13 | WT | 2019 | Nov | 0.27 | Winter | 191111 | Batch 9 |
| 1111-L1 | Δ*tssL* | 2019 | Nov | 0.21 | Winter | 191111 | Batch 9 |
| 1111-L3 | Δ*tssL* | 2019 | Nov | 0.07 | Winter | 191111 | Batch 9 |
| 1111-L5 | Δ*tssL* | 2019 | Nov | 0.08 | Winter | 191111 | Batch 9 |
| 1111-L6 | Δ*tssL* | 2019 | Nov | 0.08 | Winter | 191111 | Batch 9 |
| 1111-L7 | Δ*tssL* | 2019 | Nov | 0.14 | Winter | 191111 | Batch 9 |
| 1111-B1 | Δ*tssB* | 2019 | Nov | 0.18 | Winter | 191111 | Batch 9 |
| 1111-B2 | Δ*tssB* | 2019 | Nov | 0.09 | Winter | 191111 | Batch 9 |

### References

1. Wood, D.W., et al., The genome of the natural genetic engineer *Agrobacterium tumefaciens* C58. Science, 2001. **294**(5550): p. 2317-23.
2. Lin, J.S., L.S. Ma, and E.M. Lai, Systematic Dissection of the *Agrobacterium* Type VI Secretion System Reveals Machinery and Secreted Components for Subcomplex Formation. PLoS One, 2013. **8**(7): p. e67647.
3. Liu, A.C., et al., A citrate-inducible gene, encoding a putative tricarboxylate transporter, is downregulated by the organic solvent DMSO in *Agrobacterium tumefaciens*. J Appl Microbiol, 2008. **105**(5): p. 1372-1383.
